# Extended Similarity Methods for Efficient Data Mining in Imaging Mass Spectrometry

**DOI:** 10.1101/2023.07.27.550838

**Authors:** Nicholas R. Ellin, Ramón Alain Miranda-Quintana, Boone M. Prentice

## Abstract

Imaging mass spectrometry is a label-free imaging modality that allows for the spatial mapping of many compounds directly in tissues. In an imaging mass spectrometry experiment, a raster of the tissue surface produces a mass spectrum at each sampled *x*, *y* position, resulting in thousands of individual mass spectra, each comprising a pixel in the resulting ion images. However, efficient analysis of imaging mass spectrometry datasets can be challenging due to the hyperspectral characteristics of the data. Each spectrum contains several thousand unique compounds at discrete *m/z* values that result in unique ion images, which demands robust and efficient algorithms for searching, statistical analysis, and visualization. Some traditional post-processing techniques are fundamentally ill-equipped to dissect these types of data. For example, while principal component analysis (PCA) has long served as a useful tool for mining imaging mass spectrometry datasets to identify correlated analytes and biological regions of interest, the interpretation of the PCA scores and loadings can be non-trivial. The loadings often containing negative peaks in the PCA-derived pseudo-spectra, which are difficult to ascribe to underlying tissue biology. Herein, we have utilized extended similarity indices to streamline the interpretation of imaging mass spectrometry data. This novel workflow uses PCA as a pixel-selection method to parse out the most and least correlated pixels, which are then compared using the extended similarity indices. The extended similarity indices complement PCA by removing all non-physical artifacts and streamlining the interpretation of large volumes of IMS spectra simultaneously. The linear complexity, O(*N*), of these indices suggests that large imaging mass spectrometry datasets can be analyzed in a 1:1 scale of time and space with respect to the size of the input data. The extended similarity indices algorithmic workflow is exemplified here by identifying discrete biological regions of mouse brain tissue.

## INTRODUCTION

Imaging mass spectrometry is an important bioanalytical tool with applications in many clinical and pharmaceutical fields.^1^ In a matrix-assisted laser desorption/ionization (MALDI) imaging mass spectrometry experiment, a thinly sliced section of tissue is first mounted on a flat substrate (*i.e.*, a microscope slide), and then a chemical matrix is applied homogenously to the surface. The matrix enables efficient ionization of analytes and is selected based on the biological molecules of interest (*e.g.*, proteins, lipids, fatty acids, etc.) and the wavelength of the incident laser. A raster of the tissue with a laser produces a mass spectrum at each sampled position in a coordinate plane, resulting in hundreds-to-thousands of individual spectra that form pixels in the resulting ion images. When imaging mass spectrometry is used in an untargeted approach, each mass spectrum can contain thousands of unique compounds at discrete *m/z* values, each producing a unique ion image.^2–4^

Recent advancements improving acquisition time and throughput have resulted in substantially more spectra being acquired per imaging mass spectrometry experiment.^5–7^ Data files can contain upwards of one million spectra per image, with thousands of individual *m/z* values per spectrum, which can make data processing and analysis challenging and time-consuming for these high-dimensionality datasets.^8^ Additionally, many studies typically consist of multiple individual datasets (*i.e.*, biological and technical replicates of multiple samples), further complicating data processing and analysis.^9, 10^ Post-processing techniques such as factorization, clustering, and manifold learning are useful tools that have emerged to mine these datasets to identify biological regions of interest and better understand tissue biochemistry.^11–14^ For example, principal component analysis (PCA) has enabled the differentiation between different stages of tumor development in cancerous tissue and aided in diagnosing disease stage.^15^ Although these techniques have proven useful in elucidating molecular pathology and biochemistry in tissues, each approach comes with its own challenges and limitations.^16–19^ For example, PCA calculates the scores and loadings through linear combinations of the mean centered data. The scores are represented as spatial-expression images and the loadings are represented as pseudo-spectra. By linearly combining the *m/z* bins, the scores and loadings often result in negative values or peaks. Since negative peaks in mass spectrometry have no physical basis (*i.e.*, they would represent negative ion abundances), it can be difficult to ascribe true physical meaning to PCA results. This calls for a computational method that can examine PCA results of imaging mass spectrometry data using only physical data, such as the extended similarity indices.

Similarity measures have been applied throughout many different fields of study, to enable efficient comparisons of data.^20^ In chemistry, similarity measures have been used to compare molecules by representing specific features of their two-dimensional (2D) or three-dimensional (3D) structures as binary fingerprints. These comparisons are conducted to screen a large amount of structures in virtual databases to identify molecules that may have similar properties to a reference molecule.^21^ For example, Lavecchia *et al.* used similarity searching based on the Tanimoto similarity coefficient to discover six ligands, similar to 4-(2-Carboxybenzoyl)phthalic acid, in the NCI database that inhibited the cell division cycle 25B (Cdc25B) protein.^22^ Similarity measures have also been used for compound annotation of experimental tandem mass spectrometry (MS/MS) data.^23, 24^ MS/MS similarity calculations performing database searches by comparing experimental spectra of an unknown compound to spectra within libraries of known compounds. When analyzing complex mixtures using an untargeted technique such as liquid chromatography coupled to tandem mass spectrometry (LC-MS/MS), identifying each of the hundreds or thousands of discrete compounds in the sample can be cumbersome. With the help of similarity matching, matching the mass spectral profiles of unknown compounds to spectral libraries to facilitate identification becomes much more efficient. However, most similarity measures only compare two objects at a time, typically a reference and a test, making these measures slow and poorly scalable. Recently, we have introduced new similarity measures, called extended similarity indices, which compare multiple objects simultaneously.^25^ Instead of pairwise comparisons common to traditional similarity measures, the extended similarity indices compare an arbitrary number of objects to each other simultaneously (*i.e.*, they are *n*- ary functions). This has opened the door for new analysis techniques such as diversity picking, the study of large molecular libraries, chemical space visualization, clustering, and protein structure determination.^26–30^ The extended similarity indices provide two key advantages: they allow quantitation of the correlations between any number of objects and they can be performed with unprecedented efficiency, requiring only O(*N*) scaling.

Herein, we present a novel post-processing workflow that utilizes extended similarity-based algorithms to compare multiple mass spectra within an imaging mass spectrometry dataset. The utility of the extended similarity indices is demonstrated by comparing multiple PCA-correlated mass spectra from imaging datasets to distinguish morphological tissue regions. PCA correlated spectra within these morphological regions are expected to have more similar spectral content than non-correlated spectra from different regions. Using this proof-of-concept workflow, the extended similarity indices have shown that spectra with stronger PCA correlations also had greater similarity coefficients when they occupied morphological tissue regions. By applying the extended similarity indices, we can efficiently determine if the PCA correlated spectra truly represent physical regions of tissue through similar spectral content.

## METHODS

### Imaging Mass Spectrometry

Ten micrometer thick transverse mouse brain sections were prepared using a CM 3050S cryostat (Leica Biosystems, Buffalo Grove, IL) and stored in a -80°C freezer for storage or a desiccator for 30 minutes prior to sample preparation. A 1,5- diaminonapthalene (DAN) MALDI matrix layer was applied using a TM Sprayer (HTX Technologies, Chapel Hill, NC). Spray conditions were as follows: 10 mg/mL of DAN in 9/1 acetonitrile/water, 30°C nozzle temperature, 6 passes, 0.1 mL/min flow rate, and 25 mm track spacing. Following matrix application, samples were stored in a desiccator for 30 minutes prior to MALDI imaging mass spectrometry using a 7T solariX Fourier transform ion cyclotron resonance (FT-ICR) mass spectrometer (Bruker Daltonics, Billerica, MA) equipped with a Smartbeam II Nd:YAG laser system (355 nm, 2 kHz, 28% laser power, 100 shots per pixel). Mass spectrometry ion optics were optimized for common lipids^31^ in positive ion mode between *m/z* 400-1,000. A 256 kB time domain transient file size was used, resulting in a resolving power (full width at half maximum) of roughly 35,000 at *m/z* 760. 98% data reduction was performed during acquisition to reduce the overall file size. A 75 µm SmartWalk setting and 75 µm raster step size was used, resulting in 15,842 pixels (spectra) with a file size of 17.7 GB.

### Computational Workflow

The first step in the computational workflow is to calculate the PCA of the ions of interest (**Figure 1A**). PCA was calculated using the Scikit-learn module in Python using 22 common lipid ions (see supplemental information) with five principal components (**Figure 1B**).^32^ Pre-processing of the raw spectra was performed using root mean square (RMS) normalization and the peak maximum was selected for interval processing of the lipid ions. Next, pixel selection *(i.e*., mass spectrum selection) for similarity comparison is determined based on the results of PCA (**Figure 1C**). For pixel selection, the score values of each principal component (PC) are separated into three groups termed here: “low,” “mid,” and “high.” The low and high groups correspond to the most negative and positive PC scores, respectively, for all pixels. The mid group contains the scores closest to zero (*i.e.*, lowest in magnitude). Since each individual score value corresponds to a unique mass spectrum per PC, and each PC is orthogonal to the other PCs, the three groups of score values will occupy different tissue regions in the spatial-expression images of each PC. Hence, each low, mid, and high group of score values can be compared relative to each other to determine greater or lesser spectral similarity. In other words, if the low scores group has a greater similarity coefficient than the mid scores group, then it also has greater spectral similarity. The number of pixels selected for the low, mid, and high groups is based on the total number of pixels in the image. Therefore, a pixel percentage of “10” will select 10% of the total pixels in each PC-defined group independently. Percentages up to 33% can be used before groups start to overlap and pixels are placed in more than one of the three defined groups. As pixel selection starts at the extremes of each criterion, increasing the percentage of selected pixels increases the inclusion of pixels with scores closer to zero in the low and high groups, and the mid group will start containing more pixels with scores farther from zero. It is important to note that the same number of pixels should be selected for each group because similarity coefficients cannot be accurately compared between groups of different pixel numbers. A similarity is counted when objects have coinciding 1s in the same position of their binary fingerprints, thus the name: coincident 1s. The number of objects that have this coinciding 1 must pass a “coincidence threshold”, γ, for it to be counted as a similarity. The coincidence threshold is defined based on the number of objects being compared and has a range of [*n*mod2, *n*-1], where *n* is the total number of fingerprints, or objects.^25^ Not being able to compare similarity coefficients for low, mid, and high groups of different sizes is a result of the coincidence threshold being naturally lower for smaller groups and higher for larger groups (*viz.* smaller groups of pixels need less coincident 1s to count as a similarity while larger groups need more coincident 1s). After PCA and creating the three groups for comparison, the mass spectra are converted to binary fingerprints (**Figure 1D and 1E**).

**Figure 1.**
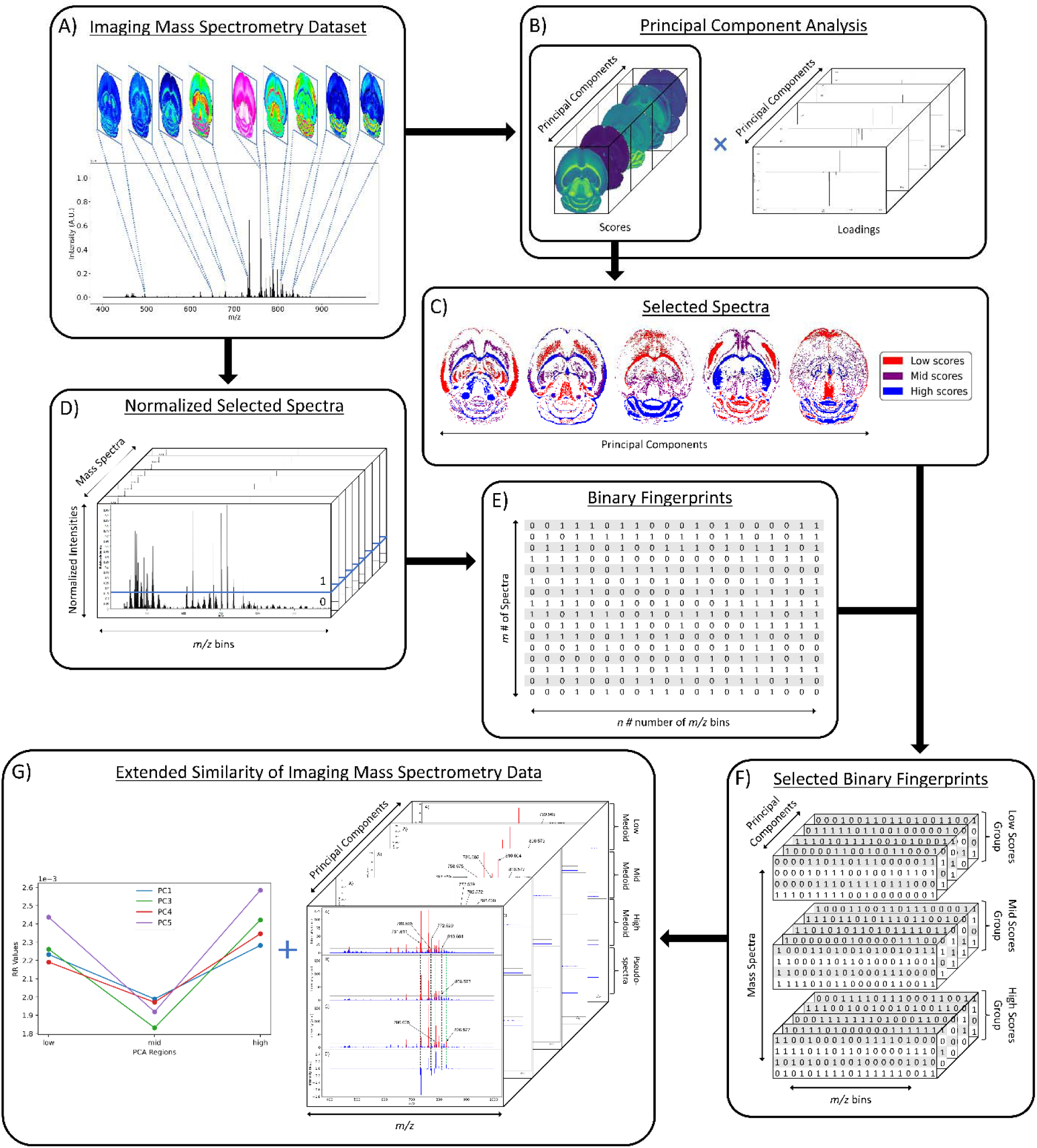
Workflow for extended similarity comparison of imaging mass spectrometry data. Overview for applying the extended similarity indices to imaging mass spectrometry data. A) Visual representation of imaging mass spectrometry data; B) PCA results of selected lipid ions; C) selected pixels based on scores for each PC; D) normalized mass spectra on 0-1 scale with intensity threshold (blue line) for binary fingerprint conversion; E) 2-D data matrix of binary fingerprints for *m* spectra and *n m/z* bins; F) selected binary fingerprints data matrix for each scores group and each PC; G) extended similarity results of each PC and respective medoid spectra.

A conversion to binary fingerprints is used to simplify the comparison framework for this proof-of-concept experiment to enable calculation of the similarity using the number of coinciding 1-bits. Future work will focus on the use of normalized real values to represent spectra rather than binary fingerprints. This conversion was performed by first extracting the raw intensity values of the imaging dataset using SCiLS Lab software (Bruker Daltonics, Billerica, MA) as a single 2D matrix of size *m* × *n*, where *m* is the number of pixels or individual spectra in the image and *n* is the number of *m/z* values (or *m/z* bins). Typical values of *m* range from 1,000-10,000 and typical values of *n* range from 250,000-500,000. However, these values can vary depending on acquisition parameters. The raw intensities are normalized on a 0-1 scale using one of four normalization methods: local, global, localTIC, or globalTIC (**Figure 1D**). Local normalization is calculated by dividing the intensity of each *m/z* bin within a spectrum by the maximum intensity in that spectrum. This process is repeated for all spectra in the image, with each spectrum being normalized to its own maximum intensity. Global normalization is calculated by dividing the intensity of each *m/z* bin within a spectrum by the maximum intensity in the entire dataset. LocalTIC normalization, or local total ion current normalization, is calculated by dividing the intensity of each *m/z* bin within a spectrum by the sum of all the intensities within that spectrum, and then repeats this process for each spectrum in the image. GlobalTIC normalization, or global total ion current normalization, is calculated by dividing the intensity of each *m/z* bin within a spectrum by the largest single total ion current pixel in the dataset. Once the spectra were normalized, an intensity threshold was defined (**Figure 1D**). The intensity threshold is a user-defined value between 0 and 1 that allows conversion to a binary format. If the normalized intensity of a peak is greater than the threshold, then it is assigned as a “1.” If the normalized intensity of a peak is less than or equal to the threshold, then it is assigned as a “0”. The result is a 2-D data matrix of 0s and 1s with *m* rows of spectra and *n* columns of *m/*z bins or bits (**Figure 1E**).

Once the spectra have been selected based on the PCA score values, normalized on a 0-1 scale, and converted to binary fingerprints, the Russel-Rao (RR) extended similarity index is calculated similar to our previous report.^25^ Briefly, for each group, all the selected binary fingerprints of the spectra are aligned into a 2D data matrix and summed together column wise. If the sum of the column is above the coincidence threshold, then it is counted as a similarity between the binary fingerprints (or spectra). A weight function is also applied to allow the columns with more coincident 1s to contribute more to the final similarity coefficient. Herein, similarity is calculated across the entire range of coincidence thresholds in 5% increments for each region within each PC.

### E-index for Comparison Workflow Parameters

Throughout this workflow, several user-defined parameters are introduced, including intensity threshold, selected pixel percentage, and coincidence threshold. Each parameter has many possible values that result in a large number of computational permutations. It is expected that there is a combination of parameters that provide the optimal distinction of similarity between the three groups of score values. To help guide the user to what the optimal set of parameters are, a new type of index, the E-index, was developed to initiate the search for the optimal intensity threshold and selected pixel percentage. The E-index identifies the optimal set of user-defined parameters by searching for the largest differences in averaged similarity coefficients between the low, mid, and high groups for each PC. Larger differences in similarity coefficients relative to the mid group result in larger E-index values. Two base functions of the E-index were developed, the maximum, *ε_p_m_* (Eqn. 1), and the robust, *ε_p_r_* (Eqn. 2).

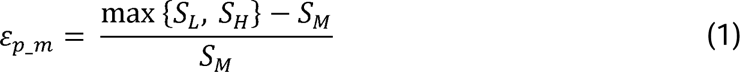

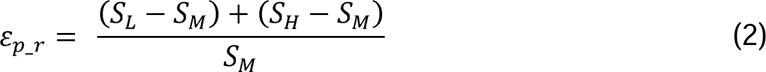

In these equations, *S* is the average similarity coefficient across all the coincidence thresholds tested, and the subscript indicates the group that the similarity value was calculated from: *L*, *M*, and *H* for the low, mid and high groups, respectively. The “max{}” function in equation 1 selects the larger of the low and high group similarities that is then subtracted by the mid similarity. The base value, ε, is calculated for a particular PC where the subscript *p* represents the respective PC (*i.e.*, one ε for every principal component, *p*). Since the base value is calculated for each PC, each combination of parameters is represented by multiple values, which is not practical. To represent each combination of parameters with a single value, a weight function is used to combine the base values, resulting in the final E-index. Three weight functions were tested to determine which will most consistently find the best set of parameters for the calculations: weighted PC (Eqn. 3), squared sum (Eqn. 4), and fraction (Eqn. 5). The weighted PC function (*E_WPC_*) will place more significance on the E-index values from the first PCs (*i.e.*, PC 1 is weighted more than PC 2 and so forth), the squared sum function (*E_Wsq_* will place more significance on the larger base values, and the fraction function (*E_Wf_*) evenly weighs the base values for each PC based on the total number of PCs calculated.

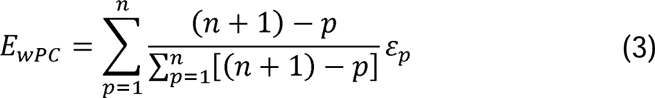

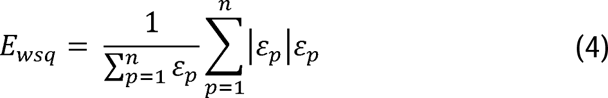

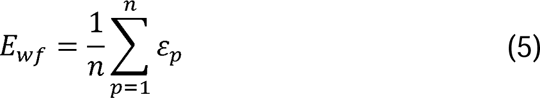

The variable *n* represents the total number of base values calculated for each combination of parameters, typically the number of PCs used to calculate the PCA. For example, if five base values were calculated, *n* would be equal to five and *p* would have values from 1-5. The variable *ε_p_* is the resulting value from the base function for every *p* principal component. To determine which combination of functions were used, some simple notation will be established. The first subscript letter after “E” will refer to which base function was used, robust or maximum, with an r or m, respectively, followed by the notations for the weight function applied, *wPC*, *wsq*, or *wf*. For example, if the robust base function was used with a squared sum weight function, the proper notation will be *E_r_Wsq_*. After running the similarity calculation for multiple intensity thresholds with selected pixel percentage values from 0-30%, the combination of base and weight functions that results in the largest E-index value should provide the best estimate for users to find the optimal set of parameters.

#### Medoid Spectra

Another application of extended similarity indices to imaging mass spectrometry data is the calculation of medoid spectra.^26^ The medoid is an object within a dataset that is the most representative point of the entire dataset. It is similar to the mean, but the medoid must be a member of the dataset. This statistical operator is ideal for MS since it takes advantage of representing the data with physically recorded values instead of averaged, non-physical points. We can apply this method to different biological regions of imaged tissue to find the most representative spectrum for that region of interest. The key challenge with current approaches to medoid calculations is that they scale as O(*N*^2^), or use approximations to go as low as O(*N*log*N*).

The medoid calculation uses the same extended similarity indices previously discussed but is iteratively applied to screen each score group to find the medoid spectrum or spectra. This is performed by removing a spectrum within a scores group and calculating the similarity coefficient of the remaining spectra without the removed spectrum (*i.e*., calculating the “complementary” similarity). The removed spectrum is then returned to the dataset, a second different spectrum is removed, and the similarity is recalculated without the other spectrum. This process is repeated for every spectrum in the dataset, with the key advantage that it can be performed in O(*N*). Once every spectrum has been iteratively removed and the similarity calculated, the iteration that resulted in the smallest similarity coefficient is identified as the medoid mass spectrum of the dataset.

The removed spectrum with the smallest complementary similarity is the medoid spectrum because it contributes most to the similarity of the spectra within the region. Since removing this spectrum resulted in the lowest similarity coefficient, it must contain the most amount of 1s that coincide with other spectra. It should be noted that this does not mean this spectrum has the most amount of 1s in its binary fingerprint. A spectrum could exist within the dataset that contains more 1s, but where none of these 1s are shared with bins in other spectra. The medoid spectra will contain the most amount of 1s that are shared with all the spectra in the dataset.

## RESULTS AND DISCUSSION

### Imaging Mass Spectrometry and Principal Component Analysis

The mouse brain lipid imaging mass spectrometry dataset acquired for this proof-of-concept study contains a total of 15,842 pixels, and each consists of 313,898 individual *m/z* bins over a *m/z* 401-1,000 mass range. Root mean square (RMS) normalization was performed on the dataset and then 22 common lipid ions were selected for PCA using five principal components. PCA aims to reduce the dimensionality of the data by explaining as much of the variance of the dataset as possible within each PC as linear combinations of *m/z* bins. This results in less variance explained with each additional PC. Therefore, earlier PCs contain the bulk of the data’s variance, ∼95%, while later PCs eventually will only contain the variance of the noise within the dataset (**Figure 2**). For this reason, PCA was calculated using only the first five principal components. The first five principal components explain 45.8229%, 34.7466%, 8.04978%, 4.51346%, and 2.24242% of the variability of the data, respectively, with a cumulative sum of 95.3751% (**Figure 2**). The spatial expression images of the PCA scores successfully differentiate multiple biological regions across all principal components (**Figure 3**). Specifically, PC 1 shows distinct separation of the cerebral cortex from the white matter of the cerebellum, midbrain, and corpus callosum (**Figure 3A**). Sub regions of the hippocampus have been observed in PC 1, 3, and 5, notably Ammon’s horn and the dentate gyrus (**Figure 3A, 3C, and 3E)**. PC 2 highlights MALDI matrix clusters derived from the DAN compound applied to the sample and does not contain any biological relevance (**Figure 3B**). PC 3 highlights the cerebellum as the major region contributing to the variability (**Figure 3C**). PC 4 visualizes the granular layer of the cerebellum, a portion of the cerebral cortex, the inferior colliculus (a sub-region of the midbrain), and the choroid plexus (**Figure 3D**). PC 5 shows many of the same regions as earlier PCs, including Ammon’s horn, the dentate gyrus, the choroid plexus, and the granular layer (**Figure 3E**). Since PCA reduces dimensionality and combines these significant structures, the pixels that contribute most to the same structures are also the most correlated (*i.e.*, the low and high groups). The pixels that contribute the least to the structures present within each PC are therefore less correlated (*i.e.*, the mid groups). The pseudo-spectra of the five principal components show the relative contribution of each ion to the variance explained by the PCs (**Figure 4**). Ions with the same sign loading are positively correlated and loadings with opposite signs are negatively correlated. A larger number of ions significantly contribute to the variance in later principal components. As a result, no single ion contributes significantly more than the others after PC 2 (**Figure 4**). As the spatial-expression image of PC 2 highlights both biological and non-biological regions (**Figure 3B**), its pseudo-spectrum correlates all lipid ions together (*i.e.*, these loadings are negative and correspond to negative scores in pixels, which are represented by the dark blue regions in the image). The only slightly positive loading (0.0109) in PC 2 is *m/z* 703.580 (**Figure 4B**), meaning this ion is very weakly expressed in the spectra that make up the positive scores (*i.e.*, the bright yellow region occupying the space just outside the tissue perimeter on the microscope slide; **Figure 3B**). This small positive correlation is likely due to a small amount of biomolecule delocalization outside of the tissue that can occur during sample preparation. Additionally, this lipid ion is more lowly abundant than the other lipids identified here (*e.g.*, roughly 10^5^ average arbitrary intensity compared to roughly 10^6^ and higher average arbitrary intensity for all other lipids analyzed).

**Figure 2.**
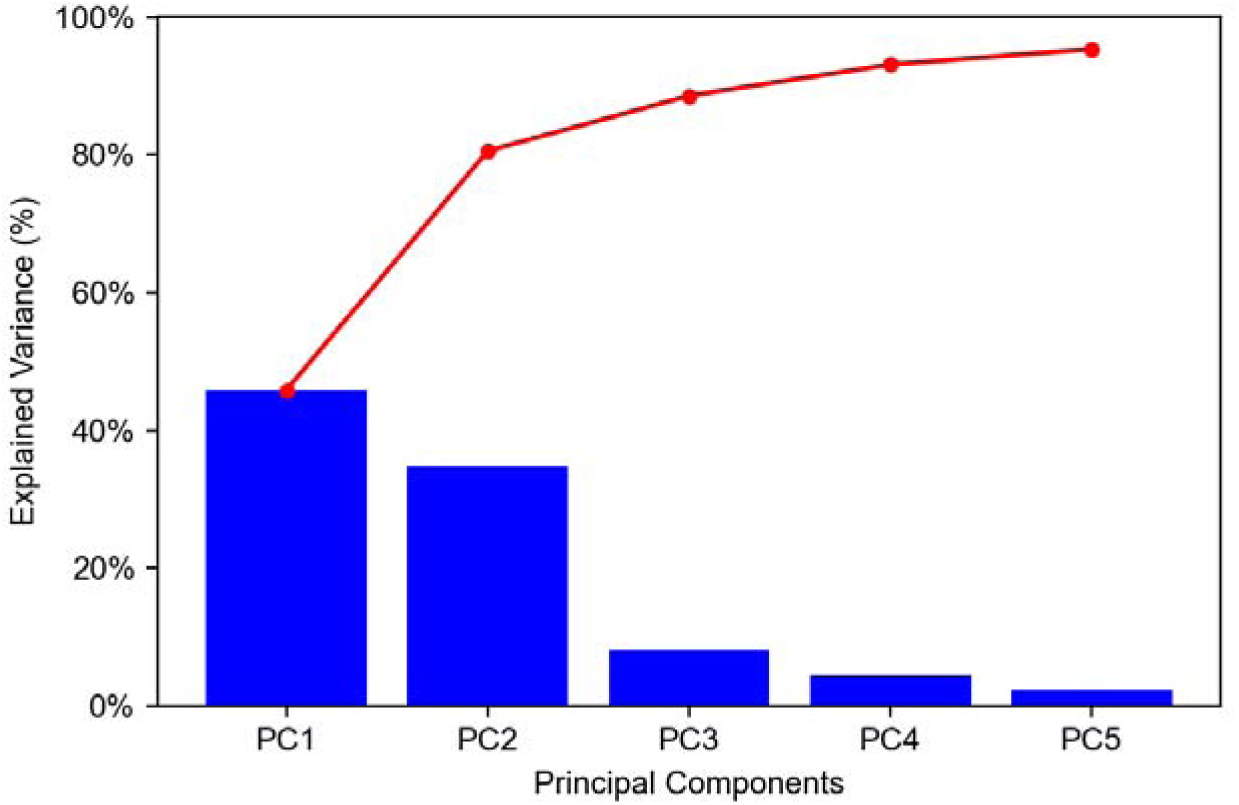
Pareto plot of explained variance for first five PCs. Explained variances for PCs 1-5 were found to be 45.8229%, 34.7466%, 8.04978%, 4.51346%, and 2.24242%, respectively. Cumulative percentages for PC 1-5 were found to be 45.8229%, 80.5695, 88.6193%, 93.1327%, and 95.3751%, respectively. Blue bars show explained variances for each PC; red trace shows cumulative explained variances.

**Figure 3.**
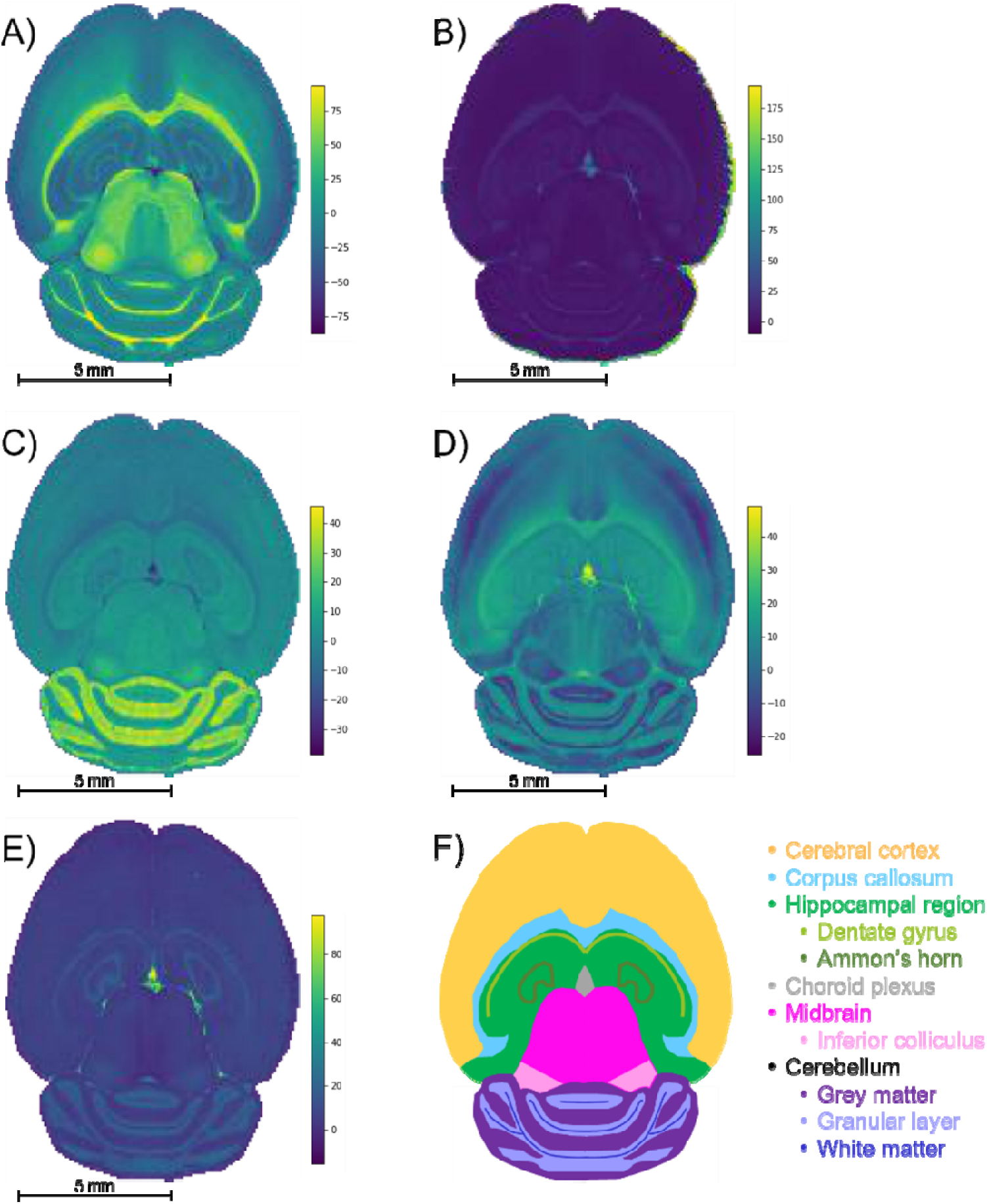
PCA spatial expression images of first five PCs. Spatial expression images for (A) PC 1, (B) PC 2, (C) principal component 3, (D) PC 4, and (E) PC 5 are compared to (F) a cartoon schematic of a mouse brain. Each color in the cartoon schematic represents a different structure of the brain (yellow, cerebral cortex; blue, corpus callosum; green, hippocampus; pink, midbrain; purple, grey matter; black, granular layer; orange, white matter). Brighter regions in the spatial expression images correspond to more positive PCA score values and darker regions correspond to more negative score values. The five PCs differentiate biological regions of the mouse brain tissue, including the corpus callosum (A and C), white matter (A), midbrain (A), Ammon’s horn (A, C, and D), dentate gyrus (A, C, and D), gray matter (C), granular layer (D), cerebral cortex (D), inferior colliculus (D), choroid plexus (C-E), and hippocampal region (E). Twenty-two lipids were chosen for dimensionality reduction. The bright regions surrounding the tissue visible in (B) correspond to non-biological matrix clusters derived from the MALDI matrix.

**Figure 4.**
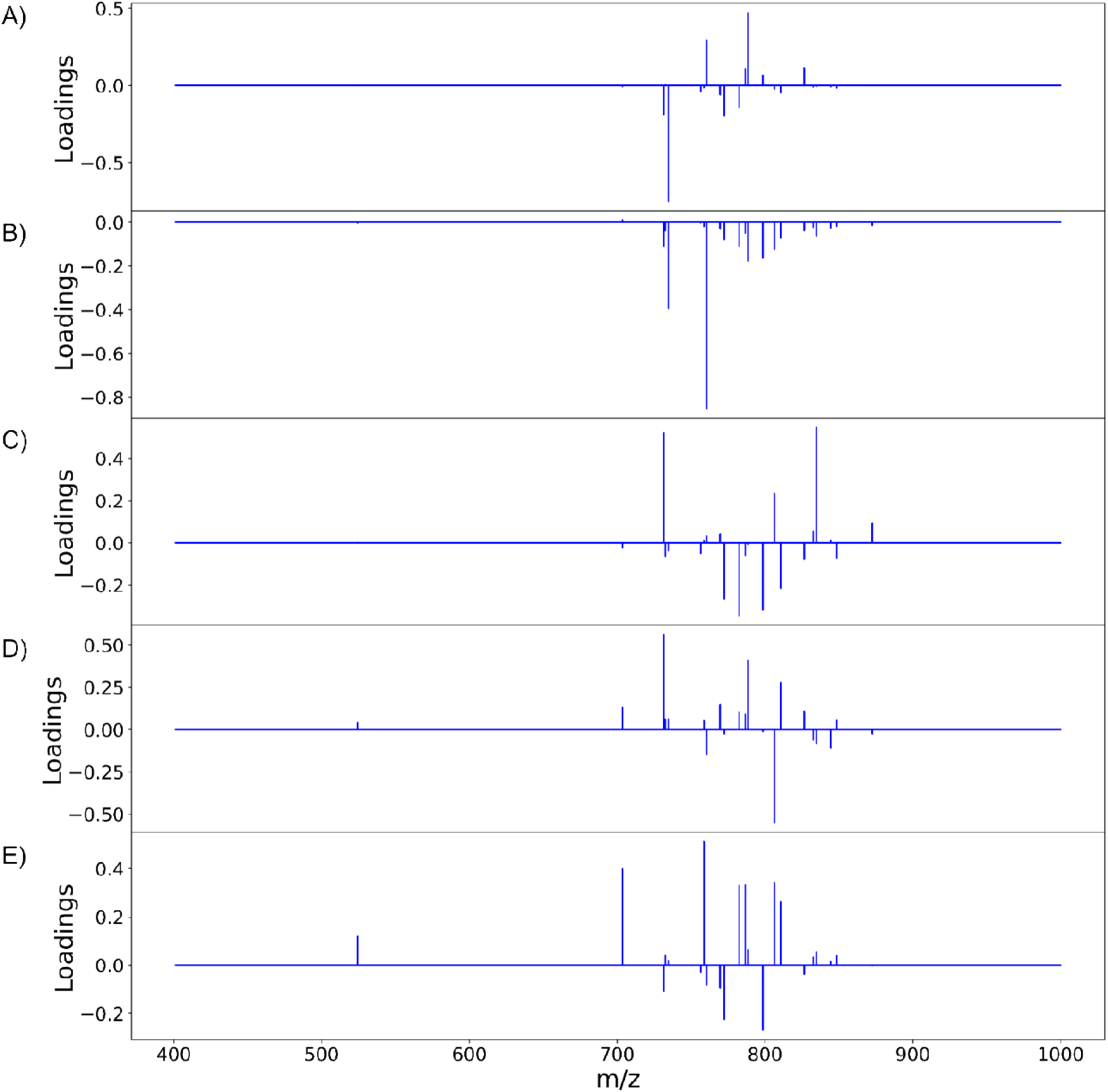
PCA pseudo-spectra of first five PCs. Pseudo-spectra are shown for (A) PC 1, (B) PC 2, (C) PC 3, (D) PC 4, and (E) PC 5. Loadings of the same sign correspond to greater positive correlation within the PC and loadings of opposite signs correspond to greater negative correlation within the PC. Twenty-one *m/z* values are contained in each pseudo-spectrum: 524.381, 703.580, 731.611, 732.558, 734.574, 756.557, 758.575, 760.588, 769.565, 772.529, 782.572, 786.605, 788.620, 798.543, 806.573, 810.604, 826.577, 832.586, 834.604, 844.528, 848.562, and 872.559. Since the peaks chosen correspond to biological regions of the mouse brain, (B) is nearly all negative because they are oppositely correlated to the non-biological matrix clusters (Figure 3B).

#### E-index

After calculation of the RR similarity coefficients with multiple sets of parameters, the E-index was used to quickly estimate the parameters that provided the largest differences in similarity between the PCA correlated regions. The two base functions (the maximum E-index, *ε_p_m_*, and the robust E-index, *ε_p_r_*) along with the three weight functions (the weighted PC function, *E_WPC_*, the squared sum function, *E_Wsq_*, and the fraction function, *E_Wf_*) for calculating the E-index were tested for each method of normalization (local, global, localTIC, or globalTIC). The same intensity thresholds and selected pixels were tested for all methods of normalization. For both “localTIC” and “globalTIC” normalization methods, the E-index values could not be calculated for most sets of parameters tested. The localTIC and globalTIC normalizations cause nearly every intensity to be below the intensity threshold due to the very large TIC value. Since nearly all of the ions are represented as 0s, many of the similarity coefficients were found to be 0 for each region rendering a similarity calculation ineffective. The E-index calculations were readily performed across all “global” and “local” normalization methods however the “local” normalization provided more consistent results across different lipid ion images and therefore will be the focus of this section (see supplemental information for other mouse brain images and normalization results).

In the mouse brain lipid dataset reported here, the first 1% of selected pixels always gave the largest E-index value for a particular intensity threshold, regardless of which combination of functions were used (**Figure 5**). In general, as the number of selected pixels for comparison increases, the E-index decreases, indicating that the difference in similarity between the score groups also decreases. The decrease is expected because as the number of pixels selected for comparison increases, their correlation through PCA weakens and thus so should spectral similarity. For some of the E-index functions, the values stabilize, causing somewhat of a plateau around 2- 10% selected pixels, where the color in the plots are relatively consistent (**Figure 5A, 5B, and 5C**). The plateaus could indicate a point where the spectra being added to the group equally express the correlations from PCA and thus differences in similarity between the score groups, E-index, are more consistent. Some plots show an increase in E-index values as the selected pixels increase, resulting in a peak at around 9% selected pixels (**Figure 5B and 5E**). When the E-index increases alongside the selected pixel percent, the mid scores group decreases in similarity while the low/high groups retain or even increase similarity as more pixels are added. The peaks seen in the *E_m_Wsq_* and *E_r_Wsq_* plots (**Figure 5B and 5E**) along with the plateaus in the *E_m_WPC_* and *E_m_Wf_* plots (**Figure 5A and 5C**) suggest the optimal selected pixels percent is within the range of 1-15%.

**Figure 5.**
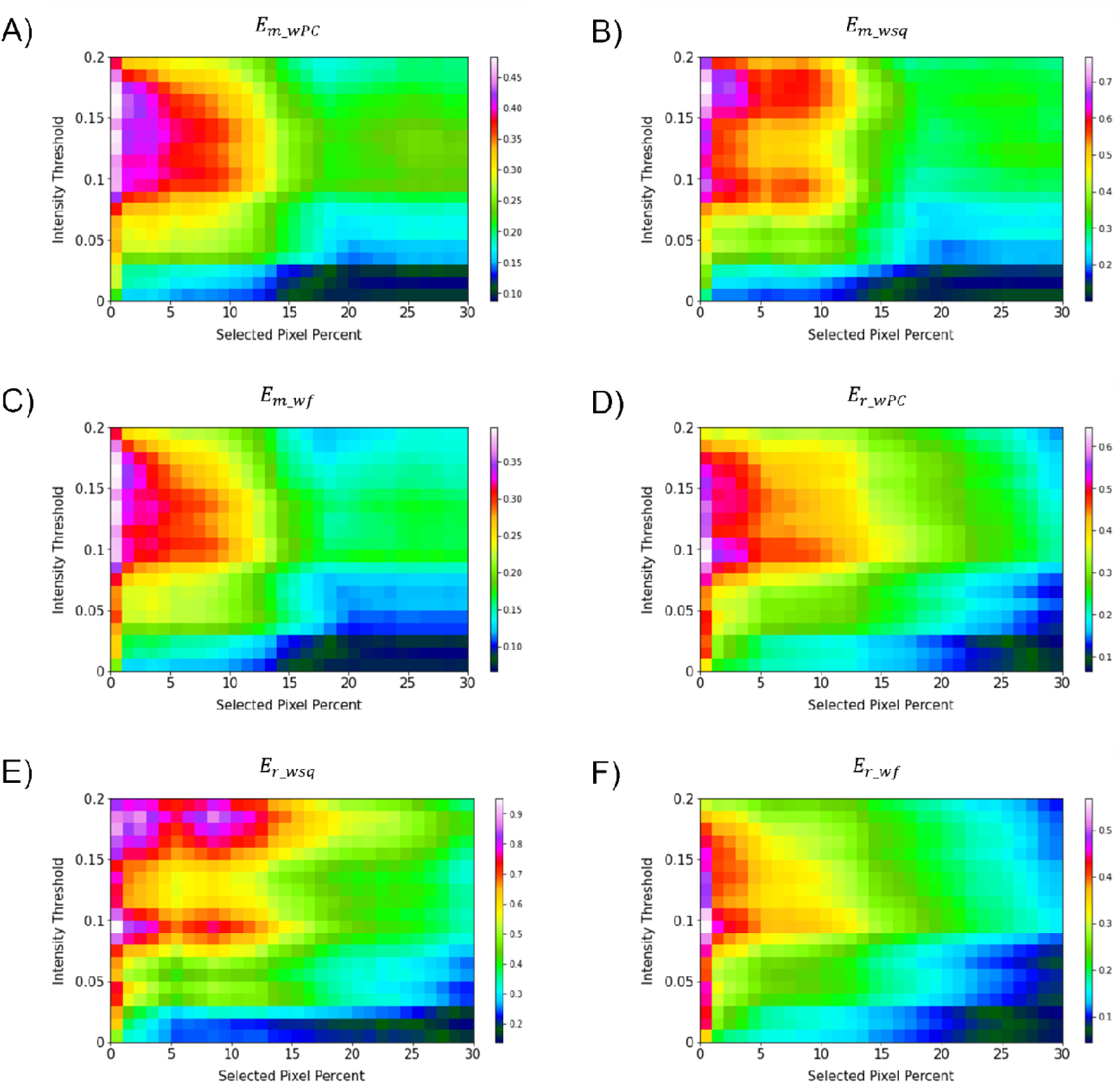
E-index plots. Two base functions (Eq 1 and 2) and three weight functions (Eqn 3-5) were tested for relative comparison. (A), (B), and (C) were calculated with the maximum base function, and weighted functions: weighted PC, weighted square, and weighted fraction, respectively. (D), (E), and (F) were calculated with the robust base function, and weighted functions: weighted PC, weighted square, and weighted fraction, respectively. Across all the methods of calculating the E-index, 1% selected pixels was found to have the largest E-index values. The optimal parameters are estimated to be within the range of 1-10% for the selected pixels and 0.09-0.19 for the intensity threshold. PC 2 was omitted from all calculations since it is highly correlated with non-biological matrix clusters.

For all pixel percentages, as the intensity threshold increases so does the E-index until about an intensity threshold of 0.09 (**Figure 5**). After an intensity threshold of 0.09, the E-index values either remain relatively constant (**Figure 5A, 5C, 5D, and 5F**) or quickly decrease and increase again creating two peaks per selected pixel percent (**Figure 5B and 5E**). As the intensity threshold increases, less *m/z* bins are assigned to 1s in the binary fingerprints, and more are assigned to 0s. The more correlated spectra from PCA will retain more of the 1s as the intensity threshold increases compared to the less correlated spectra. Since the E-index evaluates the difference in similarity between the low/high groups and the mid group, if the mid group decreases in similarity more than the low/high groups the E-index will increase. The point where the E-index plateaus is the point where increasing the intensity threshold results in the same decrease in 1s for all groups. With these trends in mind, the E-index plots point to the optimal intensity threshold being within the range of 0.09-0.19 (**Figure 5**).

The E-index serves to guide users toward the optimal set of parameters, so it is important to remember that the results provided here are not definitive. Each imaging mass spectrometry experiment will have its own range of optimal values and it is up to the user to determine them. Final determination of optimal parameters will be discussed in the following section.

#### Russel-Rao Extended Similarity Index

The RR extended similarity index is the ratio of the total number of weighted coincident 1s to the total number of bits in a single binary fingerprint. Using the RR index, the similarity is calculated based on the presence of ions above the intensity threshold, therefore providing an estimate of the spectral similarity of present ions for the group being compared. It is expected that the low and high groups exhibit a greater number of coincident 1s, or similarity, relative to the mid groups because of their stronger correlation through PCA. When the three regions follow these expected trends, the plots should create a “V” shape, meaning the low and high groups have more *m/z* bins in common within themselves than the mid groups. Therefore, using the E-index as a guide, the final evaluation of the optimal parameters for the similarity calculation was based on the presence of a “V” shape in the resulting plots (**Figures 6 and 7**). Based off this “V” shape, it was determined that an intensity threshold of 0.01 and selected pixel percentage of 1% provided the optimal distinction of similarity. The Russel-Rao extended similarity coefficients of the three regions for each PC consistently decreases as the number of coincident 1s needed to count as a similarity, or coincidence threshold, increases (**Figure 6**). This decreasing RR similarity with respect to the increasing coincidence threshold is expected because a larger coincidence threshold indicates more 1s must be shared between spectra for that bin to count as similar (**Figure 6**). The differences in RR coefficients with respect to the mid region for PCs 2 and 5 appear to decrease as the coincidence threshold is increased, meaning the distinction in similarity between the different regions becomes less significant as the coincidence threshold is increased (**Figure 6B and 6E**). Meanwhile, the opposite is seen with PCs 1, 3, and 4 (**Figure 6A, 6C, and 6D**). Additionally, the RR coefficients were found to be extremely small (10^-4^) for a 0-1 scale (**Figure 6**). The small RR coefficients are due to a large number of bins in the spectra being represented as 0s in their binary fingerprints. Since the similarity coefficients are a relative measure of similarity, small values are not necessarily indicative of a poor similarity once compared to a reference value. The mid groups were chosen to act as the reference value within each PC since they should have naturally lower similarity coefficients.

**Figure 6.**
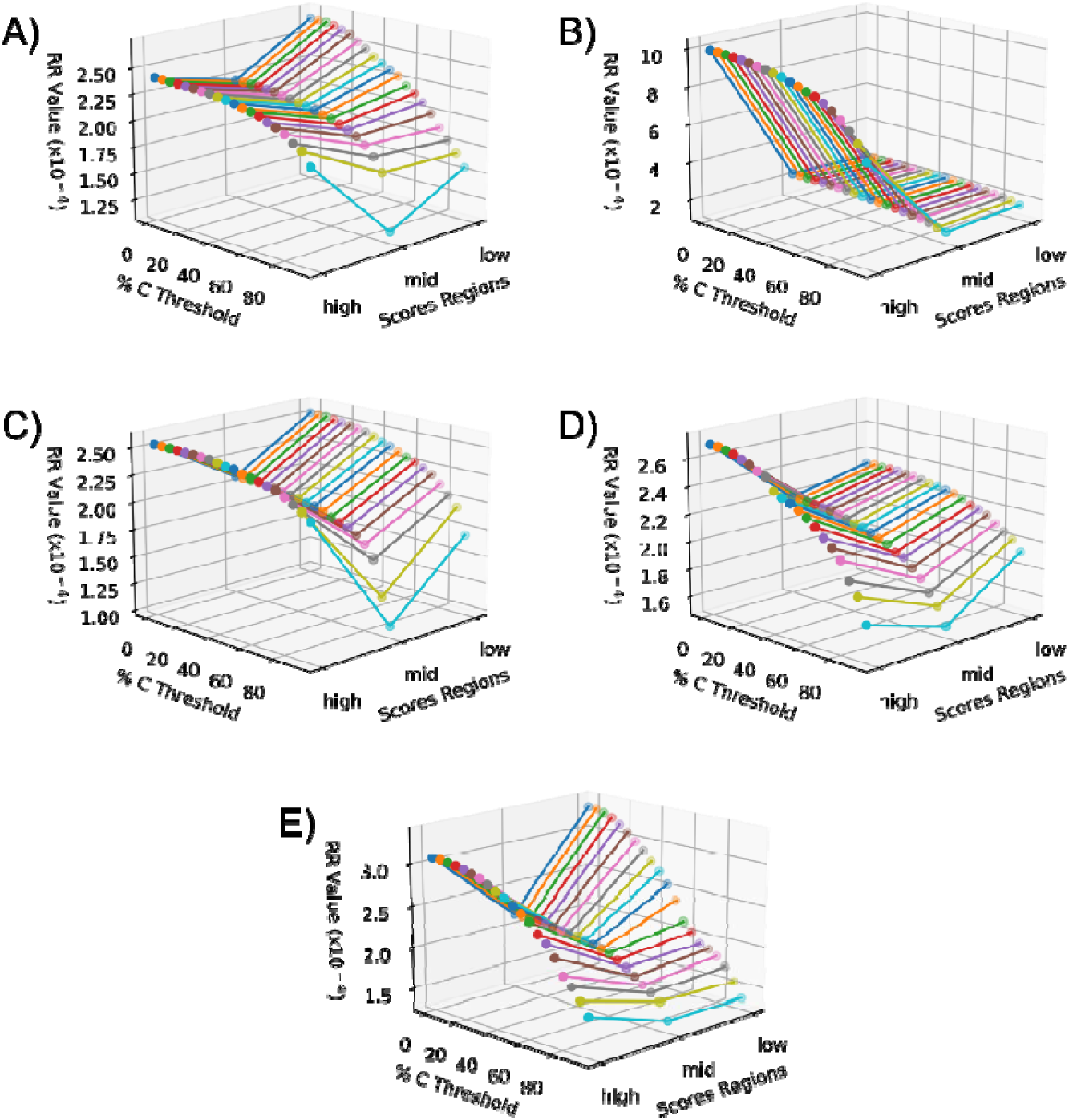
Extended similarity indices as a function of region and percent coincidence threshold for each PC. (A) PC 1, (B) PC 2, (C) PC 3, (D) PC 4, and (E) PC 5. All PCs follow the expected trend of decreasing similarity as the coincidence threshold increases due to more coincident 1s being needed to count as a similarity. The intensity threshold was set to 0.01 and the percent selected pixels was 1%. This set of parameters was determined to be the best for similarity calculation based on the better “V” shape seen in all the PCs.

**Figure 7.**
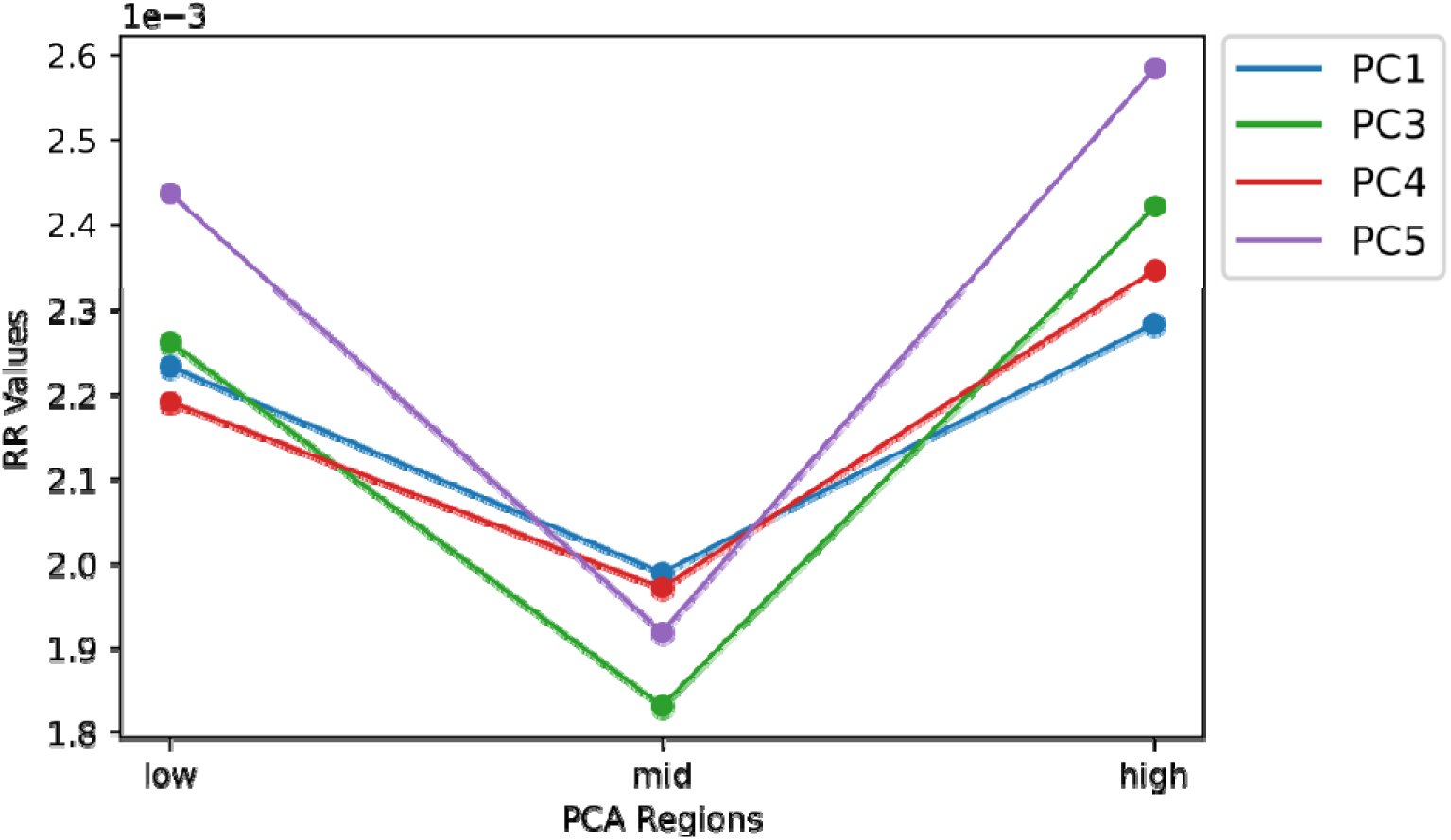
Averaged extended similarity coefficients as a function of group for each PC. Similarity Coefficients were average across the percent coincidence thresholds to represent each region for each PC with a single value. PCs 1, 3, 4, and 5 correspond to the Blue, green, red, and purple plots, respectively. Due to its non-biological correlation PC 2 was removed to provide a more accurate visualization of the data. Intensity threshold was set to 0.0 and the selected pixel percent was 1%. This set of parameters were chosen to be the optimal set since they created the best “V” shape across all PCs.

The similarity coefficients for each score group were averaged and plotted together to help compare and interpret them in a 2-dimensional plot (**Figure 7**). In this 2-dimensional plot, the differences in similarity of the three regions for each PC can be visualized. However, the similarity of the high score group of PC 2 is much greater than all the other PCs making interpretation difficult. Upon removal of PC 2, the characteristic “V” shape is observed with the remaining principal components (**Figure 7**). Looking at the spatial distribution of the score groups for each PC, the low and high groups all occupy multiple biological regions of the mouse brain tissue: cerebral cortex, white matter, gray matter, corpus callosum, hippocampus, choroid plexus, and midbrain (**Figure 8**). While the mid groups for every PC show no discernable structures (**Figure 8**), the unique spatial distribution patterns for all the score groups confirm the extended similarity indices’ ability to discern biological regions of PCA correlated spectra. The strongly correlated low and high groups occupy biological regions of tissue and have greater spectral similarity compared to their respective weakly correlated mid groups that do not occupy any biologically distinct structures (**Figures 7 and 8**). By applying the extended similarity indices to the PCA of IMS data, the correlated spectra can be efficiently connected to biological regions of tissue.

**Figure 8.**
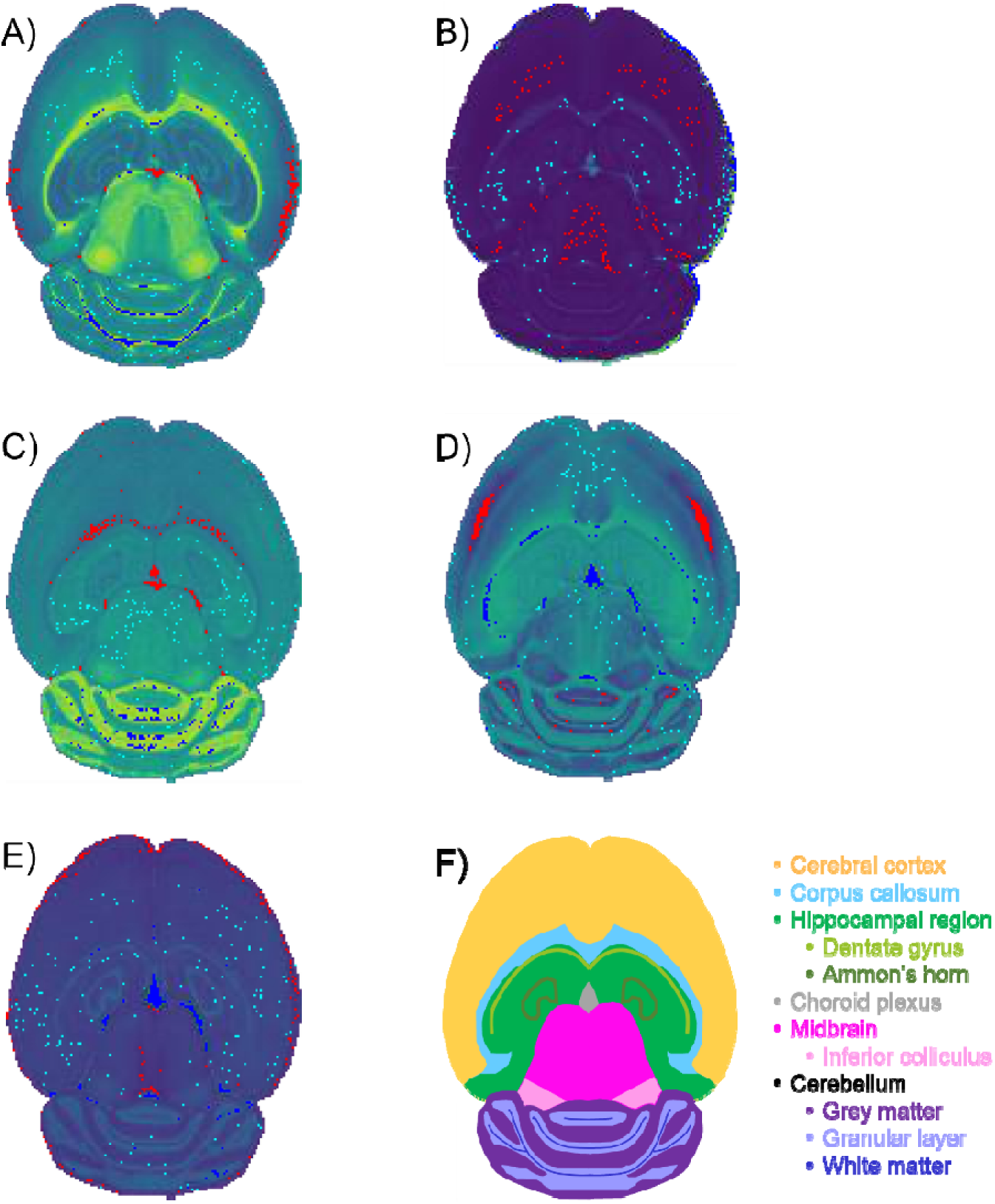
Selected pixels of optimal extended similarity indices parameters. Optimal percentage of pixels was determined to be 1% of the total pixels in the image. Red, teal, and blue colored pixels correspond to the low, mid, and high score regions each containing 1% of the total pixels in the image. The red, teal, and blue regions were overlaid with the spatial-expression images, providing the green background gradient. (A) PC 1: Red (low)- edges of cerebral cortex and choroid plexus; Teal (mid)- does not inhabit distinct structure; Blue (high)- white matter of the cerebellum. (B) PC 2: Red (low)- mostly cerebral cortex and midbrian; Teal (mid)- no distinct structures; Blue (high)- non-tissue region. (C) PC 3: Red (low)- corpus callosum and choroid plexus; Teal (mid)- no distinct structures; Blue (high)- grey matter of the cerebellum. (D) PC 4: Red (low)- cerebral cortex; Teal (mid)- no distinct structure; Blu (high)- corpus callosum and choroid plexus. (E) PC 5: Red (low)- outer most layer of the cerebral cortex; Teal (mid)- no distinct structures; Blue (high)- choroid plexus.

#### Medoid Spectra

Medoid spectra were calculated for intensity thresholds of 0.01, with 1% of the total pixels selected for each group. Only one medoid was found for each PC and each region, which is preferred since the goal is to be able to represent each region with a single spectrum. One medoid was found for each region since the lower intensity threshold allows more lowly abundant ions to be counted as 1s in the binary fingerprints and thus contribute to the similarity. If a higher intensity threshold is chosen, then many of the lowly abundant ions will never be counted as a 1 in the binary fingerprints since they innately cannot surpass the intensity threshold. On the other hand, only a few lipids were correlated to their respective region and loading (see supplemental information). For example, PC 1 had only two lipids correlated to their respective loading sign whereas 22 lipids were initially chosen for PCA. In order to connect the medoid spectra to the loadings of the PCA, the intensity threshold should be near the point of intensity variation for the lipids being analyzed. For example, highly abundant lipids will typically have a wider absolute range of ion intensities than lowly abundant lipids. The intensity threshold would need to fall within the dynamic range of the lipid intensities for it to accurately reflect the trends the PCA is trying to portray. If the intensity threshold is outside the range over which the lipid intensities vary, then the lipids will always be represented as a 1 or always represented as a 0 in every binary fingerprint. Since the E-index estimates a higher optimal intensity threshold, an intensity threshold of 0.1 was also used to calculate the medoid spectra. A value of 0.10 was chosen because a 0.1 intensity threshold and 1% selected pixels consistently provided one of the largest E-index values (**Figure 5).**

For intensity thresholds of 0.10, nearly all groups from every PC had more than one medoid spectrum. Within each group the binary fingerprints of the medoids were all nearly identical, with only a few lipids that vary between the spectra. The multiple medoid spectra that resulted for each group indicate that each spectrum contributes equally to the overall similarity of the region and equally represents the group. Ideally there should only be one medoid spectrum per group, but due to the low resolution of the binary fingerprint representation used here, more spectra can potentially have the 1s needed to be counted as a medoid. For example, each group had a unique set of lipids that were always present in the medoid’s binary fingerprint. These consistently present lipids are the most common in the group and are thus what the medoid spectra represent. As long as a spectrum contains all of these lipids in the binary fingerprint, it can be considered a medoid. For the lipids that vary between each medoid spectrum, they are not common enough within the group to count as a similarity. If one of the variable lipids within a medoid’s binary fingerprint did count as a similarity, then it would have resulted in a smaller complementary similarity and thus would need to be present in all medoid spectra.

Many of the loadings from PCA are properly correlated with their respective lipids in the binary fingerprints, such as those seen in PC 1 (**Figure 9A, 9C, and 9D**). Although highly abundant lipids are easily represented in the binary fingerprints, the lipids that are of particular interest are those that are correlated with the PCA scores and loadings. The lipids at *m/z* 786.605 and 826.577 are correlated with the positive loadings of PC 1 and are unique to the binary fingerprint of the high group medoid (**Figure 8A, 9C, and 9D**). Similarly, four lipids (*m/z* 731.611, 769.565, 772.529, and 810.604) are correlated with the negative loadings and the low group binary medoid (**Figure 8A, 9A, and 9D**). The mid group binary medoid has a mix of lipids that are correlated to both the positive and negative loadings. However, *m/z* 806.573 has a negative loading but is unique to the mid region medoid binary fingerprint (**Figure 8A, 9B, and 9D**). Since PC scores are tied to the loadings, pixels with negative scores (*i.e.,* the low group), should be directly correlated with the lipids that have negative loadings and vice versa (*e.g.*, *m/z* 810.604 and 826.577, respectively) (**Figure 9A, 9C, and 9D**).

**Figure 9.**
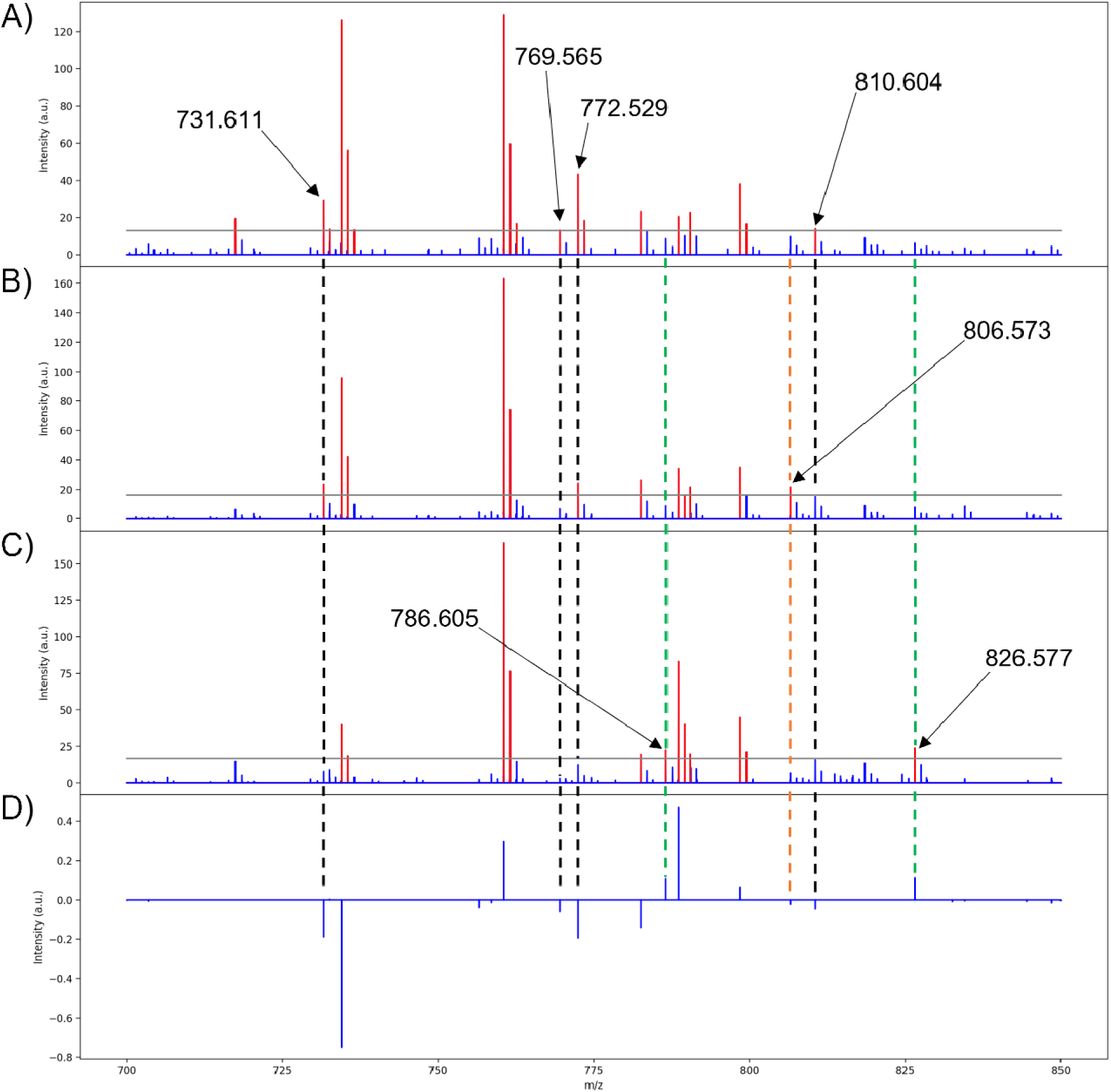
Medoid spectra of the three score groups for PC 1. Red peaks indicate intensities above the intensity threshold, blue peaks indicate intensities below the intensity threshold, and the gray line marks the intensity threshold selected for the conversion (0.10 is used here). (A) Low group medoid spectrum for PC 1, (B) mid group medoi spectrum for PC 1, (C) high group medoid spectrum for PC 1, (D) pseudo-spectrum for PC 1. Dashed lines are to show loadings that are uniquely expressed in the medoid of a PC scores group. Black, lipids with negative loadings that are only present in the low medoid binary fingerprint; green, lipids with positive loadings that are only present in the high medoid binary fingerprint; orange, lipids that are opposite of the expected trend (*i.e.*, a lipid with a positive loading but only present in the low medoid binary fingerprint and vice versa, or a lipid only present in the mid group’s medoid).

In the remaining principal components, 13 of the 22 lipids used for PCA were properly correlated to the medoids of at least one scores group and its corresponding loading when an intensity threshold of 0.10 was chosen (see supplemental information). This directly ties the extended similarity indices’ ability to help interpret PCA results by providing accurate real ion intensities for reference when analyzing the loadings and providing a real mass spectrum to represent biological regions highlighted with the score groups. Real ion intensities allow easier comparison of lipid abundances in different tissue regions (*i.e.*, a lipid that is lowly abundant throughout the entire tissue, but exhibits strong correlations, or higher loading, within one specific tissue region with other more abundant lipids). It is important to note that the mid medoid may also present the same lipids that are expected to be unique to the high and low groups, such as *m/z* 731.611 (**Figure 9A and 9B**). Although this situation might appear to indicate that the lipid is not unique to a specified group, the mid group is composed of pixels with both positive and negative score values closest to zero. Therefore, the mid group will exhibit the correlations from both low and high groups, but to a lesser extent since the magnitude of the scores is the strength in which a loading is expressed and the mid group is composed of the smallest score values. Unexpectedly, the mid group medoid could also express lipids that are absent in both the low and high groups. For example, *m/z* 806.573 is only present in the binary fingerprint of the mid group’s medoid with a loading of -0.023 (**Figure 9A, 9B, and 9D**). The expression of *m/*z 806.573 only in the mid group medoid is unexpected because scores are the strength in which a particular loading is expressed, so the low group should most strongly express all the negative loadings for that PC since it is composed of the most negative score values. Calculation of the medoid using the extended similarity indices reveals that *m/z* 806.573 does not follow the expected trend from the PCA results, further demonstrating the utility of this method in efficiently analyzing PCA data and the spatial distributions of analytes. Lipids that are present in all the medoids and binary fingerprints exist exclusively above the intensity threshold for nearly all pixels.

The medoid serves as an accurate representation of the loadings and the score groups, and thus the medoid calculation should be based on the lipids selected for PCA. For the extended similarity indices to be based on the lipids selected for PCA, the intensity threshold must be set so that the selected lipids exist above and below, or varies close to, this value. In the binary medoids with the intensity threshold set to 0.1, the properly represented loadings (*m/z* values 731.611, 769.565, 772.529, 786.605, 810.604, and 826.577) all have ion intensities that vary around the threshold value (**Figure 9A, 9C, and 9D**). The medoids calculated with the intensity threshold set to 0.01 have much fewer PCA-correlated lipids that vary around this threshold value (see supplemental information), meaning that the intensity threshold 0.01 is in the range of variance for the correlated lipids. When the intensity threshold is correctly set within the range of variance for the PCA correlated lipids, the extended similarity indices can accurately calculate a medoid to represent the correlated spectra.

## CONCLUSIONS

This proof-of-concept application of novel extended similarity indices has provided insight into the similarity of PCA-correlated mass spectral content. The extended similarity indices enable more efficient post-processing of lipid imaging mass spectrometry datasets by quickly identifying biological versus non biological tissue regions selected by PCA. The spectral content within a biological tissue region is expected to be similar, and the extended similarity indices provide a metric for determining the level of spectral similarity. Mass spectra (*i.e.*, pixels in an imaging mass spectrometry dataset) that are highly correlated by PCA and have greater similarity than the uncorrelated spectra may indicate common localization to a biological structure. Conversely, spectra that are highly correlated by PCA, but have lower spectral similarity than uncorrelated spectra, may indicate a non-biological structure resulting from linear combinations of the *m/z* bins. The extended similarity indices can also identify medoid mass spectra to represent each selected group from PCA. The medoid spectra from the selected low and high score groups of PCA were found to uniquely exhibit *m/z* values with the respective negative and positive loadings. Using real mass spectra to represent biological regions allows the non-physical loadings to be represented with real lipid ion intensities for reference when analyzing PCA results. The real lipid intensities also allow for facile comparison of relative intensities with other correlated lipids in the principal component. Knowledge of the medoid spectra and relative similarities of the three score groups can enable more definitive biological conclusions when interpreting PCA results. Future work will focus on representing spectral intensities with more precise resolution and moving away from PCA reliance by applying the extended similarity indices to other computational algorithms, such as *k*-means clustering. In order to increase the resolution of spectral intensities, the extended continuous similarity indices will be used to calculate the similarity of the spectra by representing each ion intensity with decimal values instead of binary values.^33^ The decimal values will offer more accurate representations of the ion intensities, while still retaining the efficiency and physical basis of the binary comparisons. Moving away from PCA reliance will enable the extended similarity indices to operate as an independent machine learning algorithm for imaging mass spectrometry data. The extended similarity indices’ reliance on PCA stems from its current inability to efficiently select pixels for comparison independently. By using the extended similarity indices to develop new clustering algorithms (such as novel flavors of *k*-means or density clustering), pixels can be grouped together based on spectral similarity from the beginning.^12^ These future applications of extended similarity algorithms in the computational mass spectrometry community are promising and offer a new exploratory method of mining imaging mass spectrometry data.

## ACKNOWLEDGEMENTS

This work was supported by the National Institutes of Health (NIH) under award R01 GM138660 (National Institute of General Medical Sciences [NIGMS]) and Eli Lilly and Company (BMP). NRE was supported by an Administrative Supplement for Undergraduate Research (GM138660-01S1). The Python code for the application of extended similarity indices to imaging mass spectrometry data is freely available at https://github.com/Prentice-lab-UF/Extended-Similarity-Indices-pyIMS.git.

## REFERENCES

(1) McDonnell, L. A.; Heeren, R. M. A. Imaging Mass Spectrometry. Mass Spectrom. Rev. 2007, 26 (4), 606–643. https://doi.org/10.1002/mas.20124.

(2) Nicolardi, S.; Switzar, L.; Deelder, A. M.; Palmblad, M.; van der Burgt, Y. E. M. Top-Down MALDI-In-Source Decay-FTICR Mass Spectrometry of Isotopically Resolved Proteins. Anal. Chem. 2015, 87 (6), 3429–3437. https://doi.org/10.1021/ac504708y.

(3) Bowman, A. P.; Blakney, G. T.; Hendrickson, C. L.; Ellis, S. R.; Heeren, R. M. A.; Smith, D. F. Ultra-High Mass Resolving Power, Mass Accuracy, and Dynamic Range MALDI Mass Spectrometry Imaging by 21-T FT-ICR MS. Anal. Chem. 2020, 92 (4), 3133–3142. https://doi.org/10.1021/acs.analchem.9b04768.

(4) Oras, E.; Vahur, S.; Isaksson, S.; Kaljurand, I.; Leito, I. MALDI-FT-ICR-MS for Archaeological Lipid Residue Analysis. J. Mass Spectrom. 2017, 52 (10), 689–700. https://doi.org/10.1002/jms.3974.

(5) Prentice, B. M.; Chumbley, C. W.; Caprioli, R. M. High-Speed MALDI MS/MS Imaging Mass Spectrometry Using Continuous Raster Sampling. J. Mass Spectrom. 2015, 50 (4), 703–710. https://doi.org/10.1002/jms.3579.

(6) Basu, S. S.; Regan, M. S.; Randall, E. C.; Abdelmoula, W. M.; Clark, A. R.; Gimenez-Cassina Lopez, B.; Cornett, D. S.; Haase, A.; Santagata, S.; Agar, N. Y. R. Rapid MALDI Mass Spectrometry Imaging for Surgical Pathology. *Npj Precis*. Oncol. 2019, 3 (1), 1–5. https://doi.org/10.1038/s41698-019-0089-y.

(7) Spraggins, J. M.; Djambazova, K. V.; Rivera, E. S.; Migas, L. G.; Neumann, E. K.; Fuetterer, A.; Suetering, J.; Goedecke, N.; Ly, A.; Van de Plas, R.; Caprioli, R. M. High-Performance Molecular Imaging with MALDI Trapped Ion-Mobility Time-of-Flight (TimsTOF) Mass Spectrometry. Anal. Chem. 2019, 91 (22), 14552–14560. https://doi.org/10.1021/acs.analchem.9b03612.

(8) Spraggins, J. M.; Rizzo, D. G.; Moore, J. L.; Noto, M. J.; Skaar, E. P.; Caprioli, R. M. Next-Generation Technologies for Spatial Proteomics: Integrating Ultra-High Speed MALDI-TOF and High Mass Resolution MALDI FTICR Imaging Mass Spectrometry for Protein Analysis. PROTEOMICS 2016, 16 (11–12), 1678–1689. https://doi.org/10.1002/pmic.201600003.

(9) Eberlin, L. S.; Ifa, D. R.; Wu, C.; Cooks, R. G. Three-Dimensional Vizualization of Mouse Brain by Lipid Analysis Using Ambient Ionization Mass Spectrometry. Angew. Chem. 2010, 122 (5), 885–888. https://doi.org/10.1002/ange.200906283.

(10) Takai, N.; Tanaka, Y.; Inazawa, K.; Saji, H. Quantitative Analysis of Pharmaceutical Drug Distribution in Multiple Organs by Imaging Mass Spectrometry. Rapid Commun. Mass Spectrom. 2012, 26 (13), 1549–1556. https://doi.org/10.1002/rcm.6256.

(11) Hjartarson, D. Extension of Maximum Autocorrelation Factorization: With Application to Imaging Mass Spectrometry Data. 2019.

(12) Verbeeck, N.; Caprioli, R. M.; Van de Plas, R. Unsupervised Machine Learning for Exploratory Data Analysis in Imaging Mass Spectrometry. Mass Spectrom. Rev. 2020, 39 (3), 245–291. https://doi.org/10.1002/mas.21602.

(13) McCombie, G.; Staab, D.; Stoeckli, M.; Knochenmuss, R. Spatial and Spectral Correlations in MALDI Mass Spectrometry Images by Clustering and Multivariate Analysis. Anal. Chem. 2005, 77 (19), 6118–6124. https://doi.org/10.1021/ac051081q.

(14) Abdelmoula, W. M.; Balluff, B.; Englert, S.; Dijkstra, J.; Reinders, M. J. T.; Walch, A.; McDonnell, L. A.; Lelieveldt, B. P. F. Data-Driven Identification of Prognostic Tumor Subpopulations Using Spatially Mapped t-SNE of Mass Spectrometry Imaging Data. Proc. Natl. Acad. Sci. 2016, 113 (43), 12244–12249. https://doi.org/10.1073/pnas.1510227113.

(15) Cho, Y.-T.; Chiang, Y.-Y.; Shiea, J.; Hou, M.-F. Combining MALDI-TOF and Molecular Imaging with Principal Component Analysis for Biomarker Discovery and Clinical Diagnosis of Cancer. Genomic Med. Biomark. Health Sci. 2012, 4 (1), 3–6. https://doi.org/10.1016/j.gmbhs.2012.04.022.

(16) Dill, A. L.; Eberlin, L. S.; Costa, A. B.; Zheng, C.; Ifa, D. R.; Cheng, L.; Masterson, T. A.; Koch, M. O.; Vitek, O.; Cooks, R. G. Multivariate Statistical Identification of Human Bladder Carcinomas Using Ambient Ionization Imaging Mass Spectrometry. Chem. – Eur. J. 2011, 17 (10), 2897–2902. https://doi.org/10.1002/chem.201001692.

(17) Hu, H.; Yin, R.; Brown, H. M.; Laskin, J. Spatial Segmentation of Mass Spectrometry Imaging Data by Combining Multivariate Clustering and Univariate Thresholding. Anal. Chem. 2021, 93 (7), 3477–3485. https://doi.org/10.1021/acs.analchem.0c04798.

(18) Wehrli, P. M.; Michno, W.; Blennow, K.; Zetterberg, H.; Hanrieder, J. Chemometric Strategies for Sensitive Annotation and Validation of Anatomical Regions of Interest in Complex Imaging Mass Spectrometry Data. J. Am. Soc. Mass Spectrom. 2019, 30 (11), 2278–2288. https://doi.org/10.1007/s13361-019-02327-y.

(19) Alexandrov, T.; Kobarg, J. H. Efficient Spatial Segmentation of Large Imaging Mass Spectrometry Datasets with Spatially Aware Clustering. Bioinformatics 2011, 27 (13), i230–i238. https://doi.org/10.1093/bioinformatics/btr246.

(20) Choi, S.-S.; Cha, S.-H.; Tappert, C. C. A Survey of Binary Similarity and Distance Measures. 6.

(21) Todeschini, R.; Consonni, V.; Xiang, H.; Holliday, J.; Buscema, M.; Willett, P. Similarity Coefficients for Binary Chemoinformatics Data: Overview and Extended Comparison Using Simulated and Real Data Sets. J. Chem. Inf. Model. 2012, 52 (11), 2884–2901. https://doi.org/10.1021/ci300261r.

(22) Lavecchia, A.; Di Giovanni, C.; Pesapane, A.; Montuori, N.; Ragno, P.; Martucci, N. M.; Masullo, M.; De Vendittis, E.; Novellino, E. Discovery of New Inhibitors of Cdc25B Dual Specificity Phosphatases by Structure-Based Virtual Screening. J. Med. Chem. 2012, 55 (9), 4142–4158. https://doi.org/10.1021/jm201624h.

(23) Bittremieux, W.; Schmid, R.; Huber, F.; van der Hooft, J. J. J.; Wang, M.; Dorrestein, P. C. Comparison of Cosine, Modified Cosine, and Neutral Loss Based Spectrum Alignment For Discovery of Structurally Related Molecules. J. Am. Soc. Mass Spectrom. 2022, 33 (9), 1733–1744. https://doi.org/10.1021/jasms.2c00153.

(24) Li, Y.; Kind, T.; Folz, J.; Vaniya, A.; Mehta, S. S.; Fiehn, O. Spectral Entropy Outperforms MS/MS Dot Product Similarity for Small-Molecule Compound Identification. Nat. Methods 2021, 18 (12), 1524–1531. https://doi.org/10.1038/s41592-021-01331-z.

(25) Miranda-Quintana, R. A.; Bajusz, D.; Rácz, A.; Héberger, K. Extended Similarity Indices: The Benefits of Comparing More than Two Objects Simultaneously. Part 1: Theory and Characteristics†. J. Cheminformatics 2021, 13 (1), 32. https://doi.org/10.1186/s13321-021-00505-3.

(26) Miranda-Quintana, R. A.; Rácz, A.; Bajusz, D.; Héberger, K. Extended Similarity Indices: The Benefits of Comparing More than Two Objects Simultaneously. Part 2: Speed, Consistency, Diversity Selection. J. Cheminformatics 2021, 13 (1), 33. https://doi.org/10.1186/s13321-021-00504-4.

(27) Rácz, A.; Mihalovits, L. M.; Bajusz, D.; Héberger, K.; Miranda-Quintana, R. A. Molecular Dynamics Simulations and Diversity Selection by Extended Continuous Similarity Indices. J. Chem. Inf. Model. 2022, 62 (14), 3415–3425. https://doi.org/10.1021/acs.jcim.2c00433.

(28) Flores-Padilla, E. A.; Juárez-Mercado, K. E.; Naveja, J. J.; Kim, T. D.; Alain Miranda-Quintana, R.; Medina-Franco, J. L. Chemoinformatic Characterization of Synthetic Screening Libraries Focused on Epigenetic Targets. Mol. Inform. 2022, 41 (6), 2100285. https://doi.org/10.1002/minf.202100285.

(29) Dunn, T. B.; Seabra, G. M.; Kim, T. D.; Juárez-Mercado, K. E.; Li, C.; Medina-Franco, J. L.; Miranda-Quintana, R. A. Diversity and Chemical Library Networks of Large Data Sets. J. Chem. Inf. Model. 2022, 62 (9), 2186–2201. https://doi.org/10.1021/acs.jcim.1c01013.

(30) Chang, L.; Perez, A.; Alain Miranda-Quintana, R. Improving the Analysis of Biological Ensembles through Extended Similarity Measures. Phys. Chem. Chem. Phys. 2022, 24 (1), 444–451. https://doi.org/10.1039/D1CP04019G.

(31) Zemski Berry, K. A.; Hankin, J. A.; Barkley, R. M.; Spraggins, J. M.; Caprioli, R. M.; Murphy, R. C. MALDI Imaging of Lipid Biochemistry in Tissues by Mass Spectrometry. Chem. Rev. 2011, 111 (10), 6491–6512. https://doi.org/10.1021/cr200280p.

(32) Pedregosa, F.; Varoquaux, G.; Gramfort, A.; Michel, V.; Thirion, B.; Grisel, O.; Blondel, M.; Prettenhofer, P.; Weiss, R.; Dubourg, V.; Vanderplas, J.; Passos, A.; Cournapeau, D. Scikit-Learn: Machine Learning in Python. Mach. Learn. PYTHON.

(33) Rácz, A.; Dunn, T. B.; Bajusz, D.; Kim, T. D.; Miranda-Quintana, R. A.; Héberger, K. Extended Continuous Similarity Indices: Theory and Application for QSAR Descriptor Selection. J. Comput. Aided Mol. Des. 2022, 36 (3), 157–173. https://doi.org/10.1007/s10822-022-00444-7.

