## Supplementary Information for "Extended Similarity Methods for Efficient Data Mining in Imaging Mass Spectrometry"

Extra IMS datasets and extended similarity results

All Python scripts used to calculate indices and generate plots can be found at: https://github.com/Prentice-lab-UF/Extended-Similarity-Indices-pyIMS.git.

For extra data not shown here, visit respective links for each data set:

https://github.com/Prentice-lab-UF/Extended-Similarity-Indices-mouse-brain-image-publication-supplemental-data

https://github.com/Prentice-lab-UF/Extended-Similarity-Indices-mouse-brain-image-A

https://github.com/Prentice-lab-UF/Extended-Similarity-Indices-mouse-brain-image-B

https://github.com/Prentice-lab-UF/Extended-Similarity-Indices-mouse-brain-image-C

Lipids for Principal Component Analysis

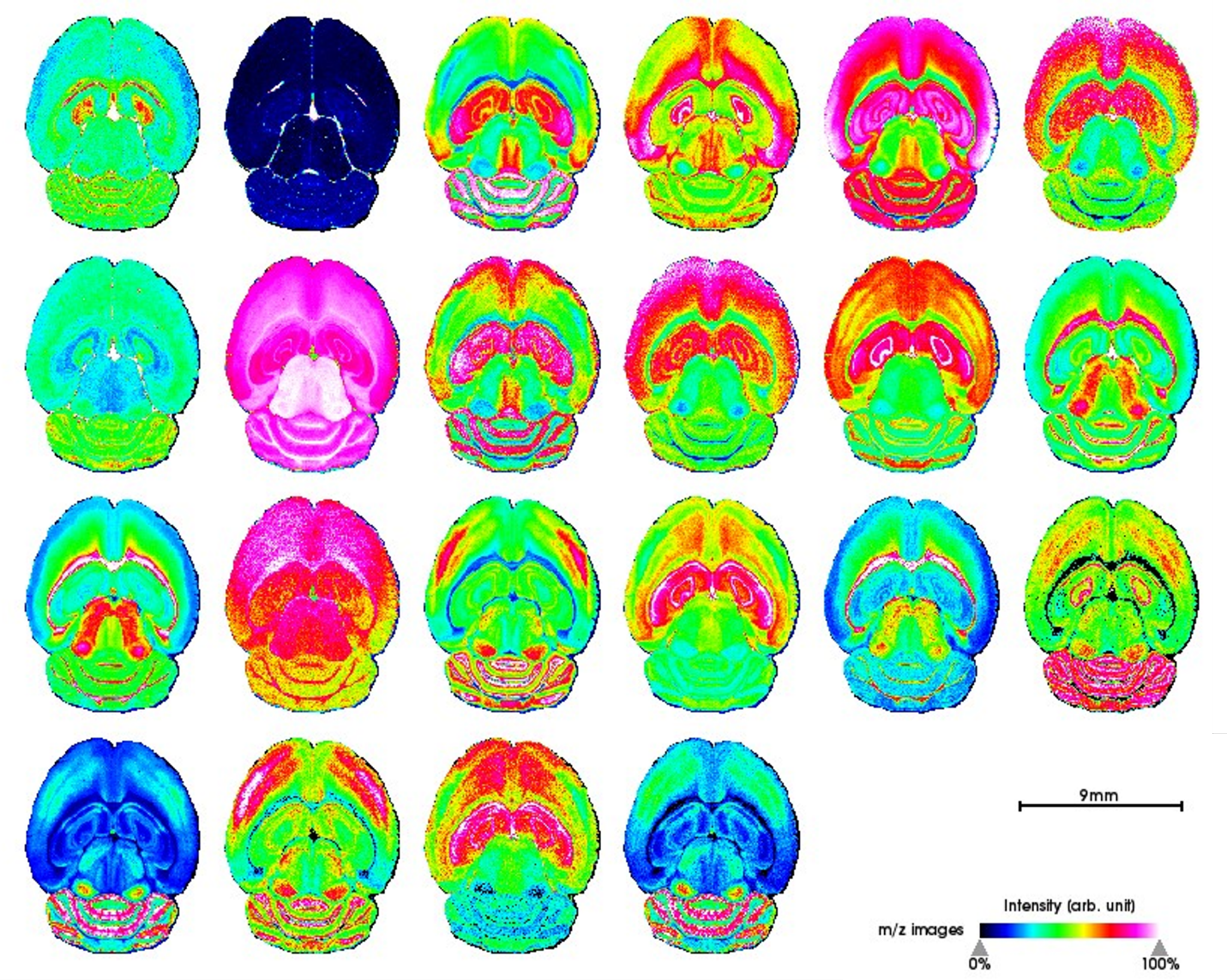

Root mean square normalized ion images for used for principal component analysis. From top left to bottom right *m/z* values are as follows: 524.381, 703.580, 731.611, 732.558, 734.674, 756.5571, 758.575, 760.588, 769.565, 772.529, 782.572, 786.605, 788.620, 798.543, 806.573, 810.604, 826.577, 832.586, 834.604, 844.528, 848.562, and 872.559.

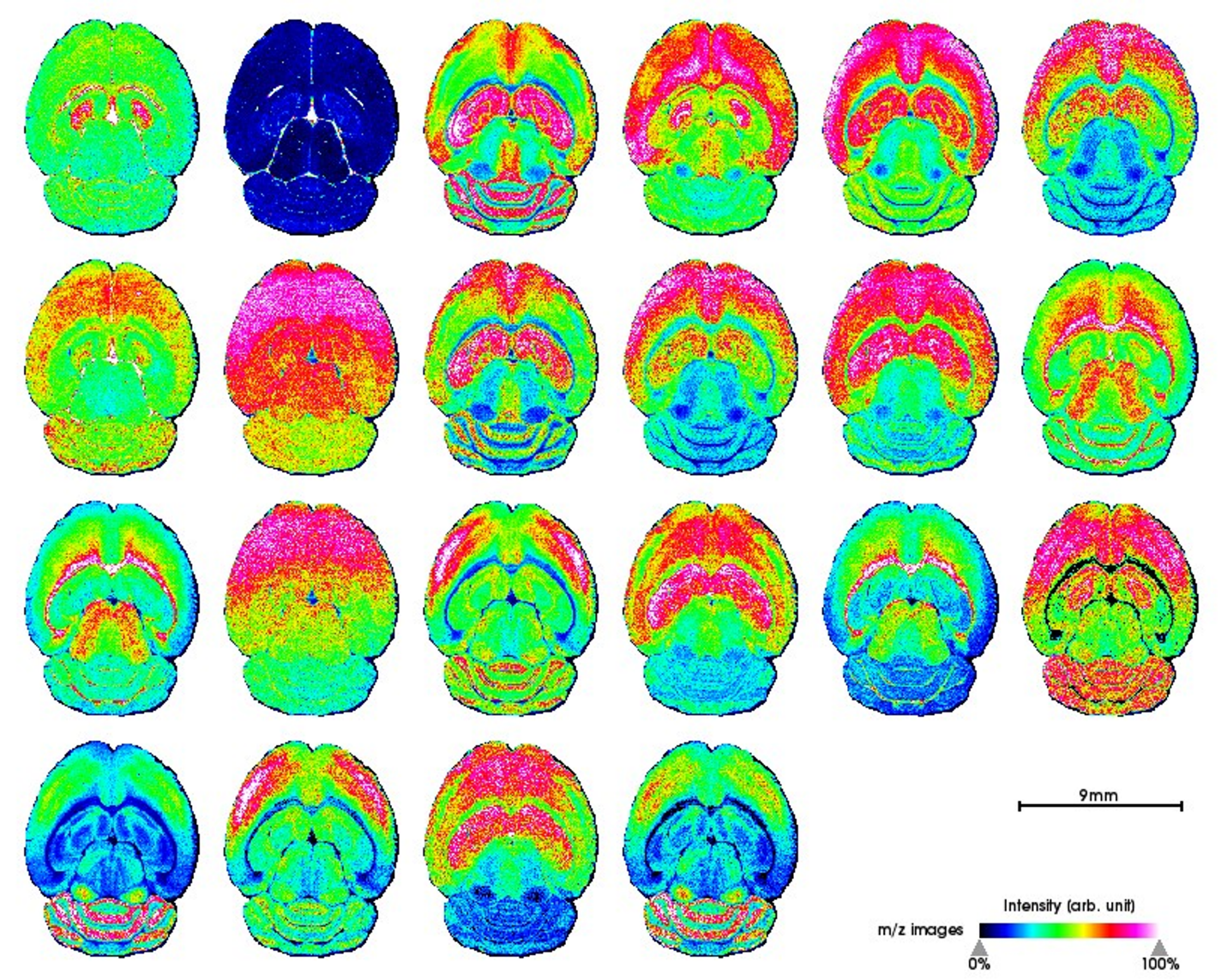

Ion images for used for principal component analysis. From top left to bottom right *m/z* values are as follows: 524.381, 703.580, 731.611, 732.558, 734.674, 756.5571, 758.575, 760.588, 769.565, 772.529, 782.572, 786.605, 788.620, 798.543, 806.573, 810.604, 826.577, 832.586, 834.604, 844.528, 848.562, and 872.559.

Positive Ion Phospholipid Identification by High Mass Accuracy

| Experimentally measured | Calculated*^a^* | ppm | Assignment |
| --- | --- | --- | --- |
| 524.3812 | 524.3711 | 19.26116828 | PC(18:0/OH) |
| 703.5759 | 703.5748 | 1.563444285 | SM(d18:1/16:0) |
| 731.6106 | 731.6061 | 6.150850847 | SM(d18:1/18:0) |
| 734.5737 | 734.5694 | 5.853769569 | PC(16:0/16:0) |
| 756.5571 | 756.5514 | 7.534187367 | PC(16:0/16:0)+Na |
| 758.5755 | 758.5694 | 8.041452766 | PC(34:2) |
| 760.5884 | 760.5851 | 4.338764985 | PC(34:1) |
| 769.5654 | 769.5620 | 4.418097567 | SM(d18:1/18:0)+K |
| 772.5295 | 772.5253 | 5.436715147 | PC(32:0)+K |
| 782.5715 | 782.5670 | 5.750306364 | PC(34:1)+Na |
|  | 782.5694 | 2.683468073 | PC(36:4) |
| 786.6054 | 786.6007 | 5.975077317 | PC(36:2) |
| 788.6197 | 788.6164 | 4.184543968 | PC(36:1) |
| 798.5434 | 798.5410 | 3.005481246 | PC(34:1)+K |
| 806.5728 | 806.5670 | 7.190971116 | PC(36:3)+Na |
|  | 806.5694 | 4.215384318 | PC(38:6) |
| 810.6042 | 810.5983 | 7.278574357 | PC(36:1)+Na |
|  | 810.6007 | 4.317785563 | PC(38:4) |
| 826.5767 | 826.5723 | 5.323188304 | PC(36:1)+K |
| 832.5857 | 832.5827 | 3.603245659 | PC(38:4)+Na |
| 834.6041 | 834.6007 | 4.073804395 | PC(40:6) |
| 844.5279 | 844.5253 | 3.078652587 | PC(38:6)+K |
| 848.5617 | 848.5566 | 6.010206037 | PC(38:4)+K |
| 872.5591 | 872.5566 | 2.865143648 | PC(40:6)+ |

a) *Calculated using "The LIPID MAPS® Lipidomics Gateway, https://www.lipidmaps.org/"*

Non-normalized Principal Component Analysis

PCA was calculated with Python module sklearn. SVD_solver was set to randomized with iterated_power of 10000 and 5 components.

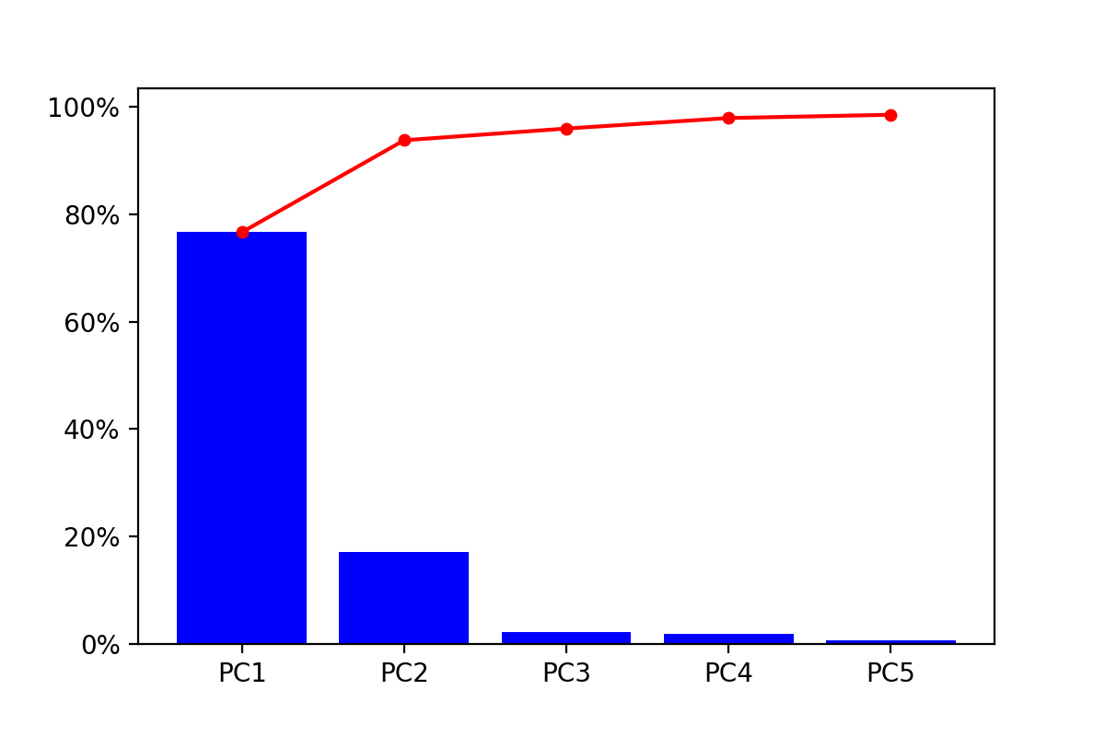

Pareto plot of the explained variance for non-normalized PCA. Explained variance for PC 1-5 are as follows: 76.7%, 17.1%, 2.18%, 1.96%, and 0.606%. The cumulative sum of the variance explained are as follows: 76.6%, 93.8%, 96.0, 97.9%, and 98.5%.

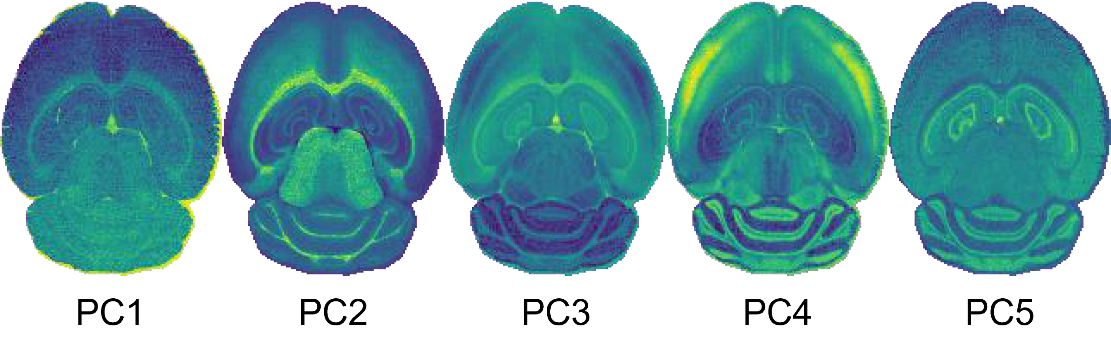

Spatial expression images for PC 1-5. Biological images similar to what was found in the RMS pre-processed PCA spatial-expression images can be seen.

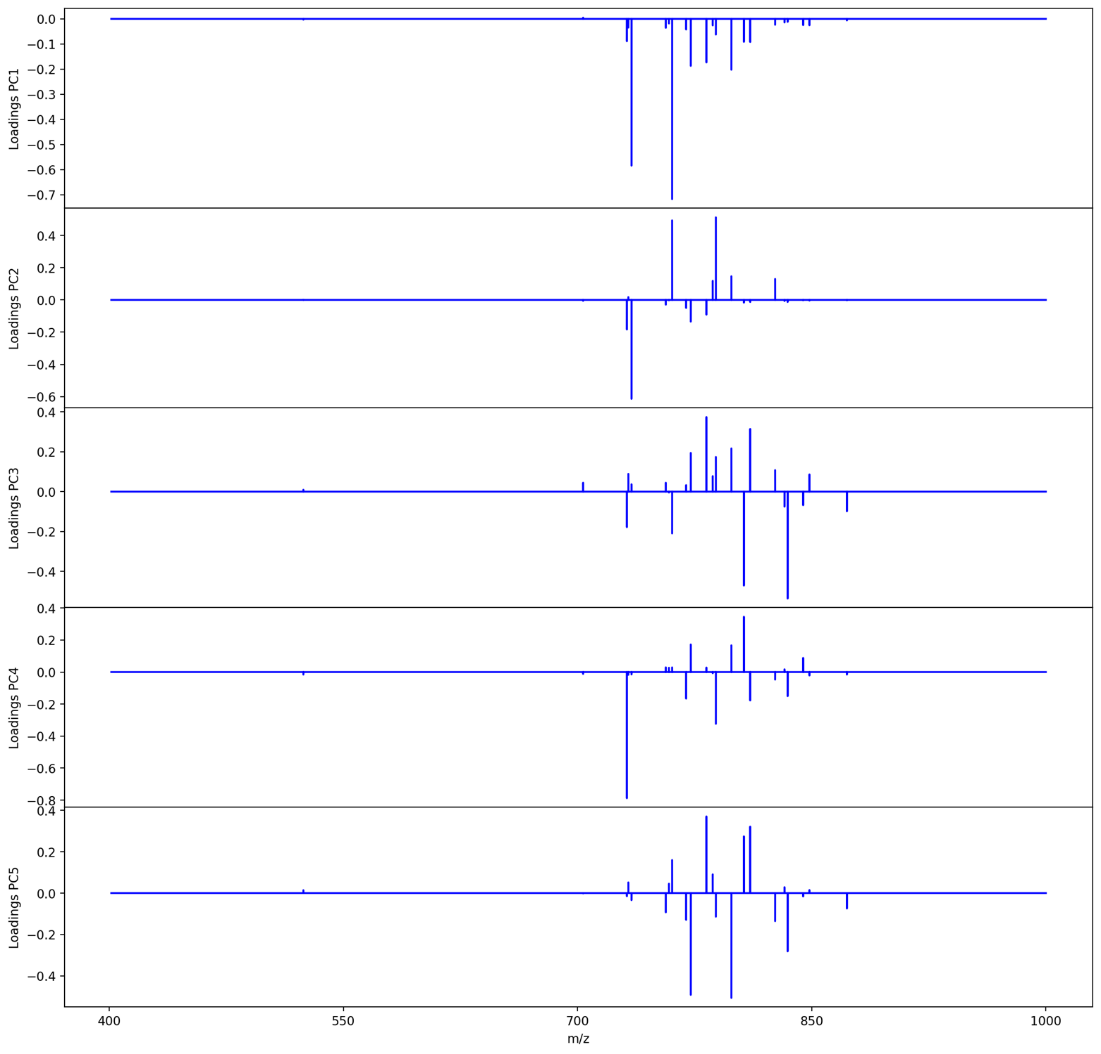

Pseudo-spectra for PC 1-5 from top to bottom.

Normalization Methods for Binary Fingerprint Conversion

Each panel contains the same mass spectra from three unique pixels in the imaging mass spectrometry dataset that are normalized using A) local normalization, B) global, C) localTIC normalization, and D) globalTIC normalization. The colored peak(s) in each spectrum highlight the peak(s) used for normalization. *I_m/z_* is used to denote the raw intensity, *i_0-1_* is used to denote the resulting normalized intensity on a 0-1 scale, *­I_max_* is to denote the largest intensity within the mass spectrum of the corresponding color, and ∑ indicates the mass range of summed ion intensities.

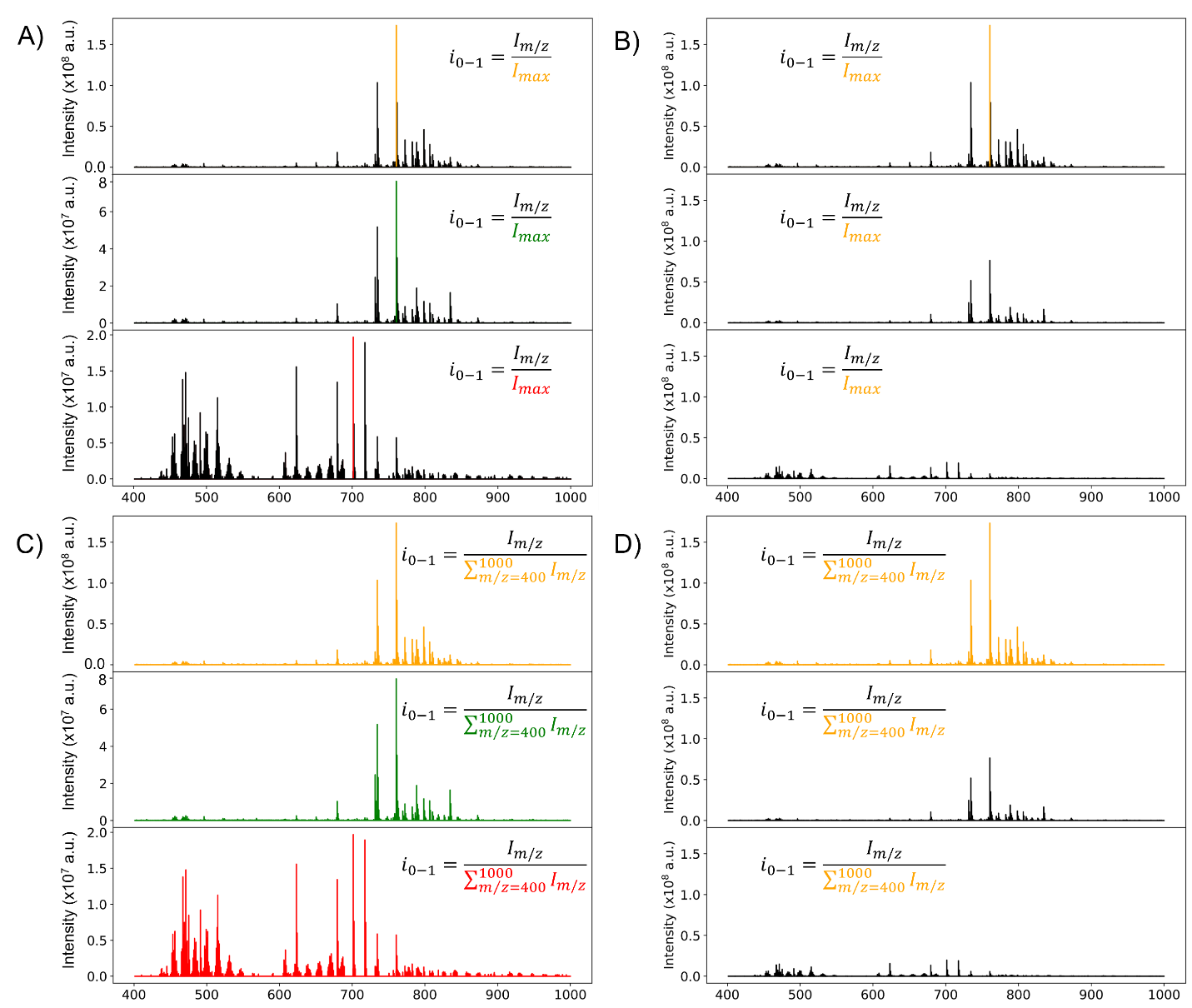

E-index Plots

PC 1 was omitted from the E-index calculations from non-normalized PCA, since it corresponds to non-tissue regions. Negative E values would indicate that the mid region has greater similarity, meaning the normalization method could not properly represent the spectra and the extended similarity calculations failed.

E-index could not be calculated for globalTIC normalized data as the intensities were suppressed too low to be above the threshold and coincide across multiple spectra.

global normalization from non-normalized PCA results

The E-index fails to show any relevant trends that could point to an optimal set of parameters. Both robust and max functions with the squared weight show flat lines with spiked in positive and negative directions. The spikes do not correspond to optimal parameters.

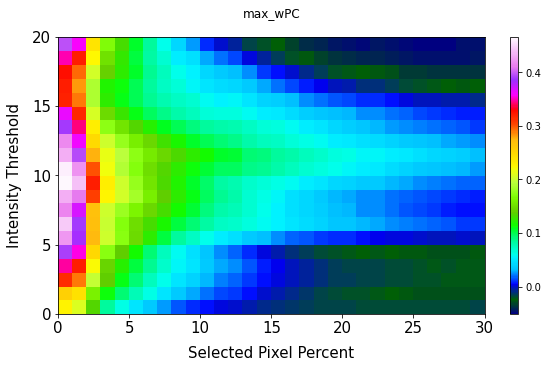

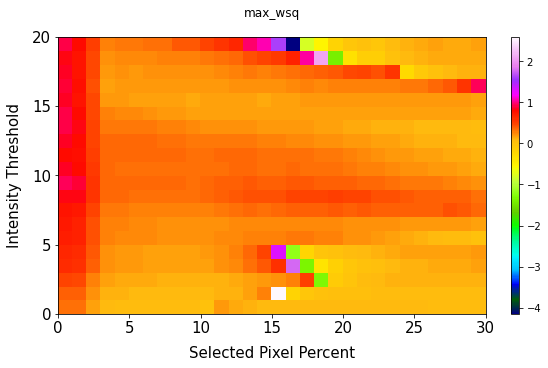

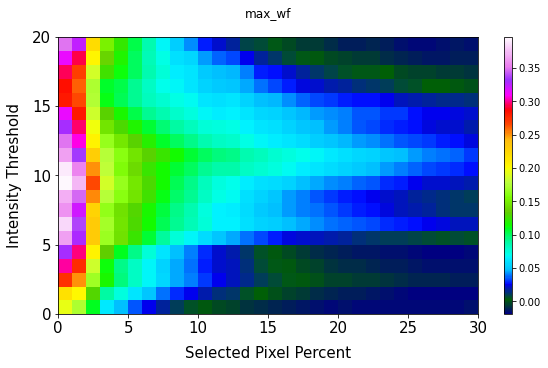

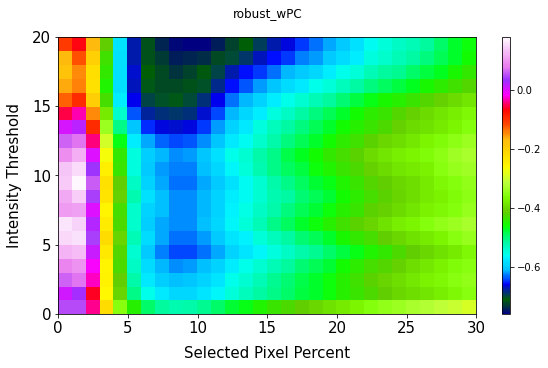

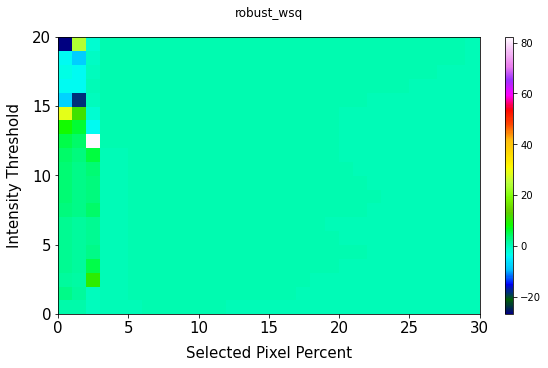

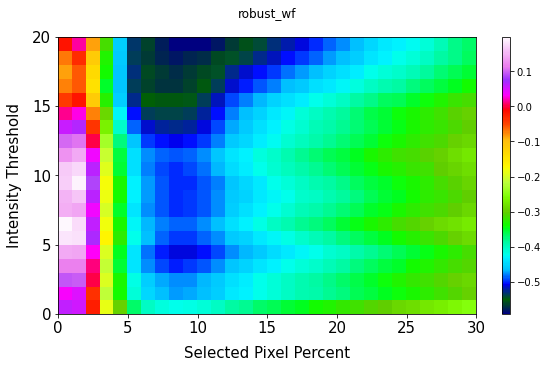

Local normalization from non-normalized PCA results

The E-index fails to show any relevant trends that could point to an optimal set of parameters. The max function with the squared weight show flat lines with spiked in positive and negative directions. The spikes do not correspond to optimal parameters. A gradual decrease in E-index values does point to local normalization be a better method of correlating the spectra.

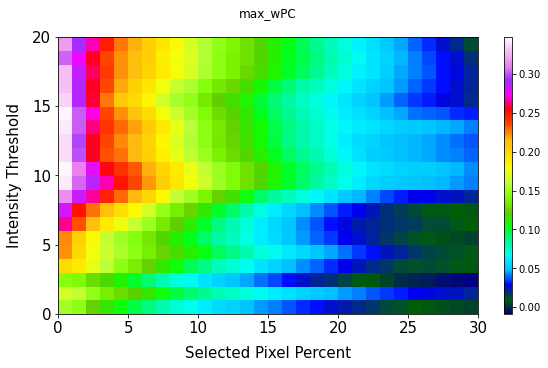

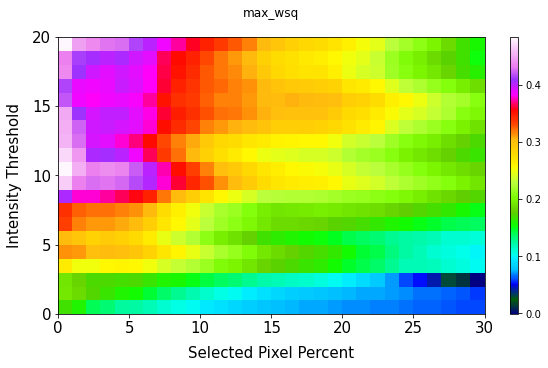

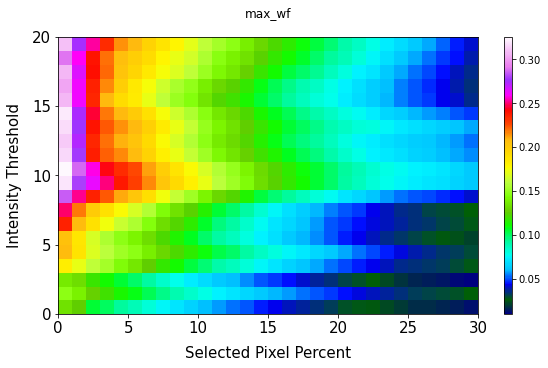

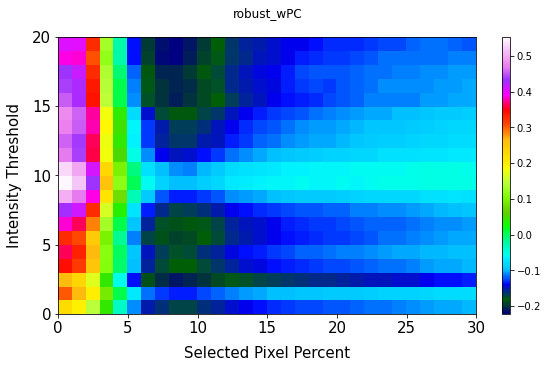

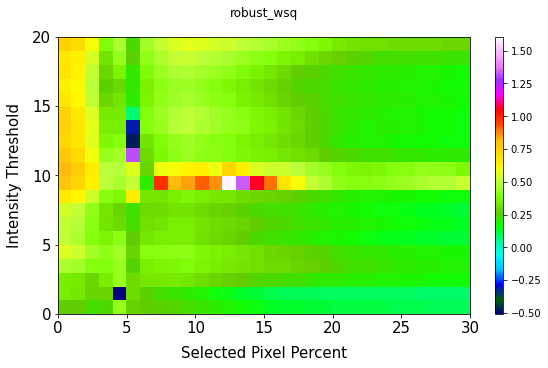

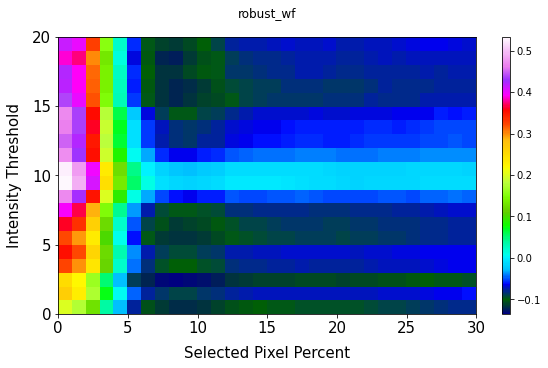

Global normalization from root mean squared normalized PCA results

Here we see the expected trend of E-values where we see greater values of E in the first 10% of selected pixels and lower values for larger amounts of pixels.
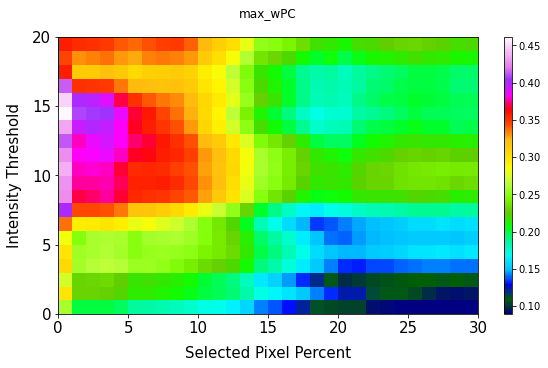

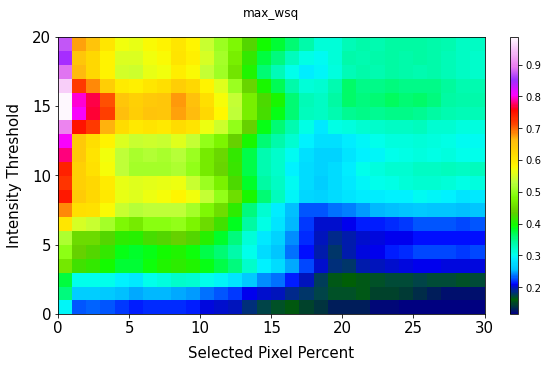

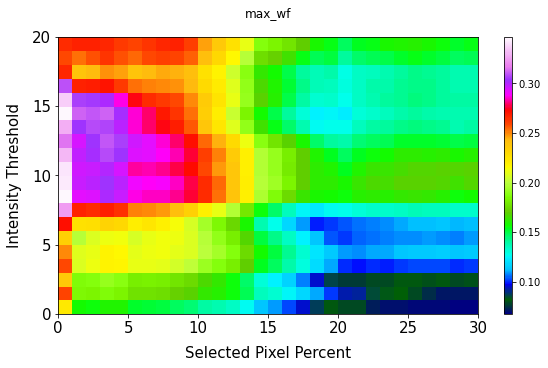

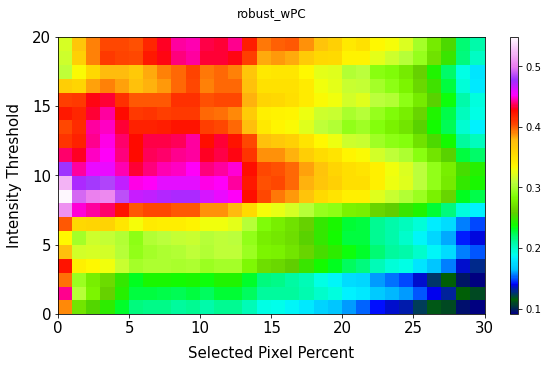

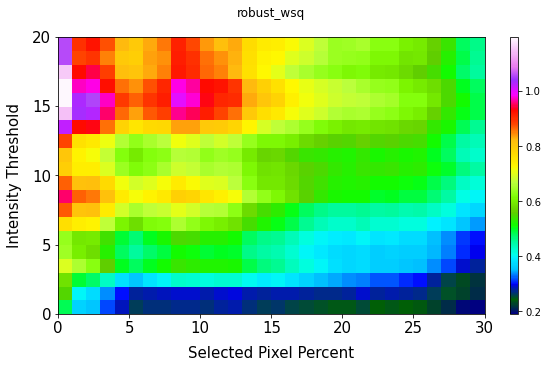

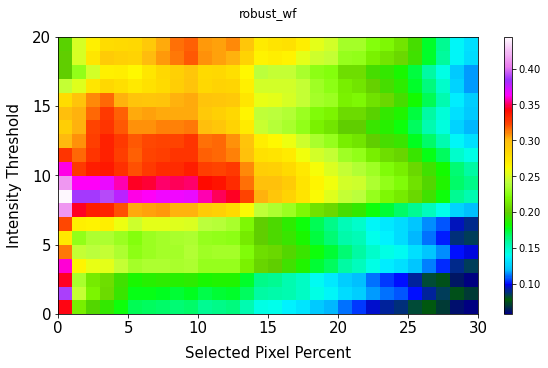

Russel-Rao 2-D V Plots

2-D V plots for local normalization with root mean squared normalized results

RMS normalized PCA was used for pixel selection. Left column has the 2D V plots with PC 2. Right column has the 2D V plots without PC 2. Distinct “V” shape can be seen up to ~20% for intensity threshold 0.01. However, the “V” shape begins to diminish after 14%. This corresponds to the decrease in E values that looks to find these large distinctions in similarity between the regions.

For intensity threshold 0.10 the V shape is not as apparent but still present for PC 3 and disappears for other PCs sooner.

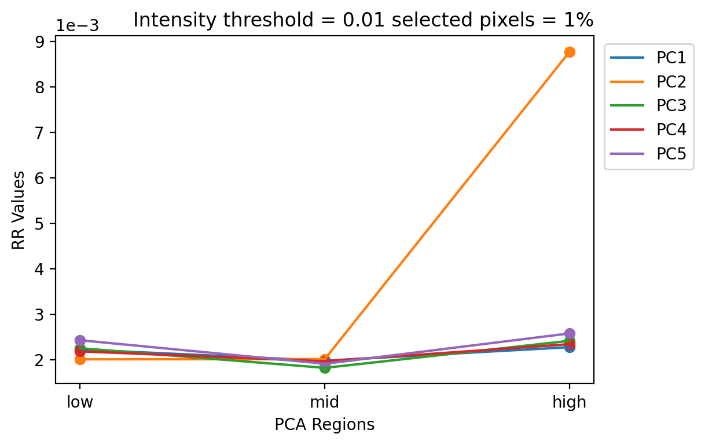

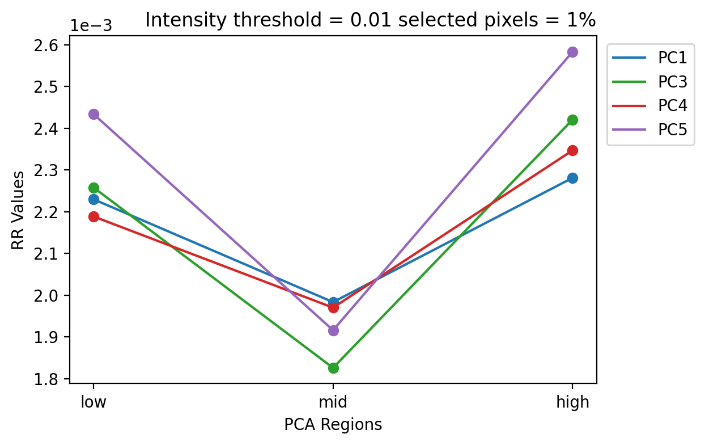

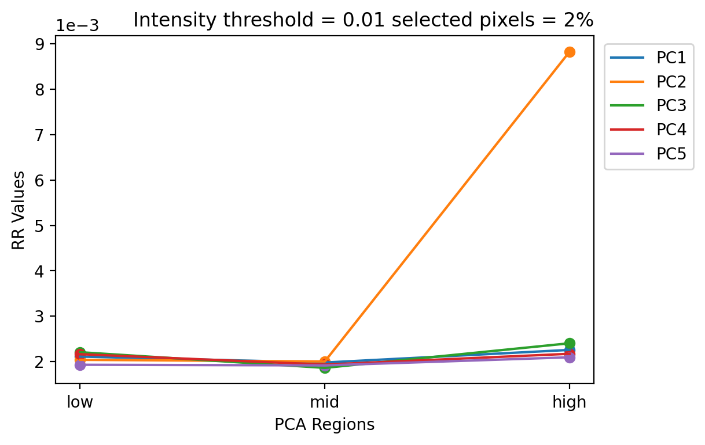

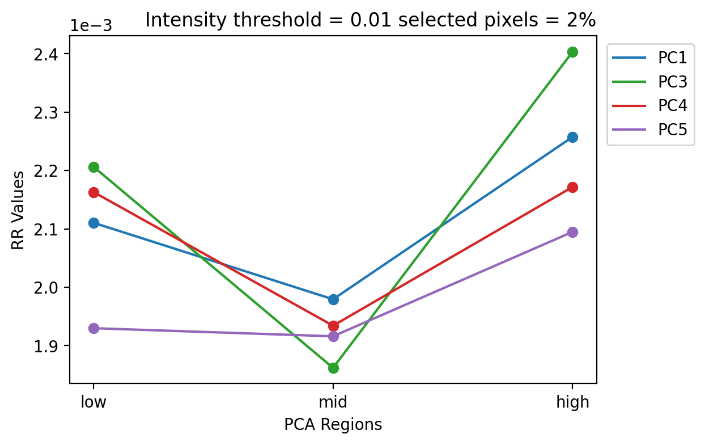

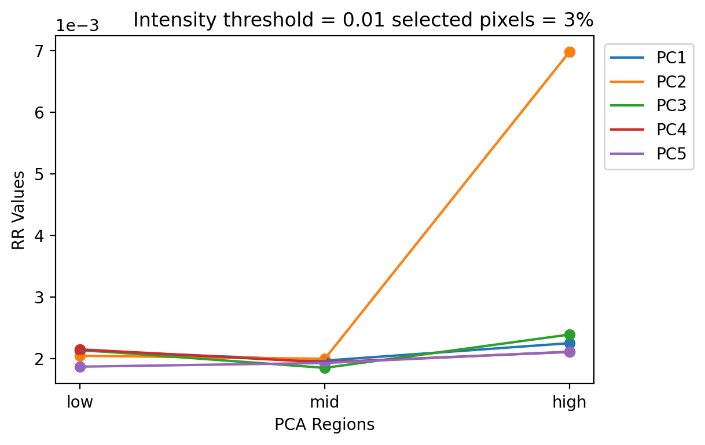

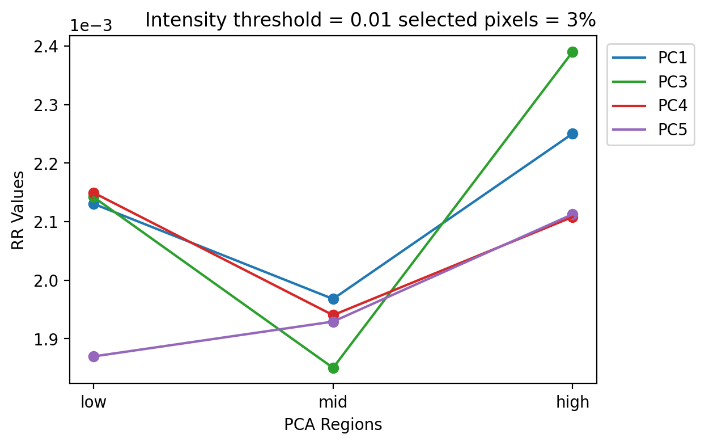

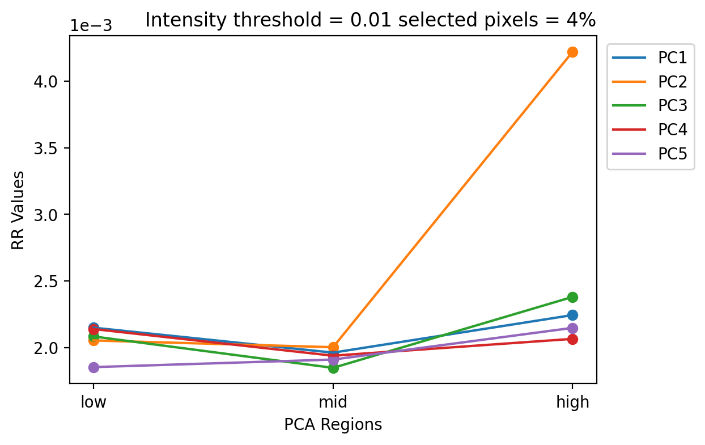

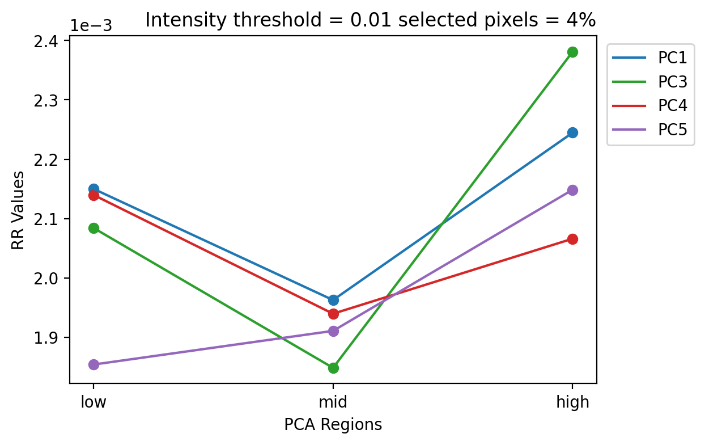

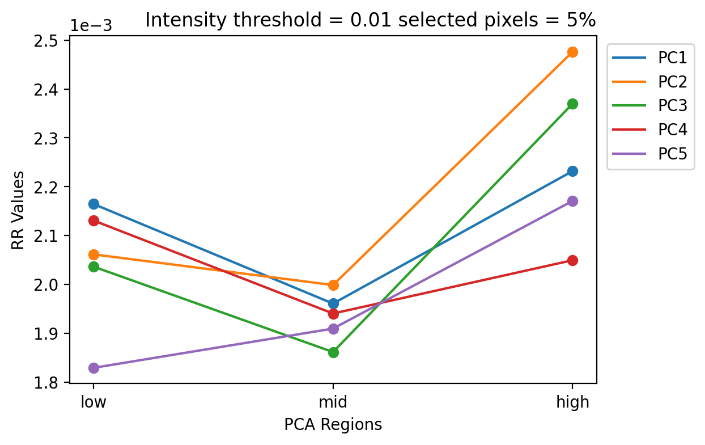

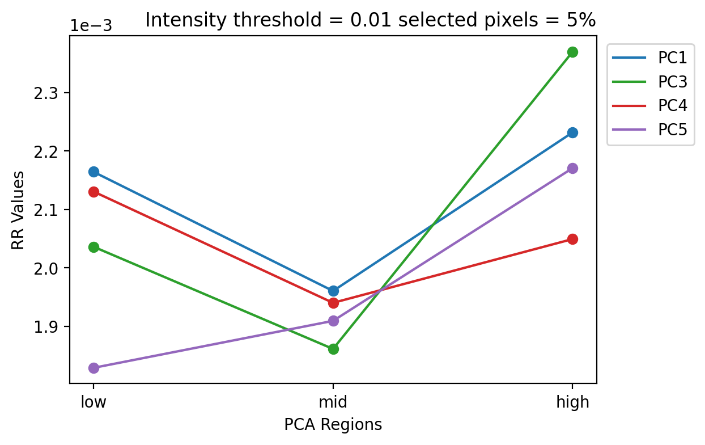

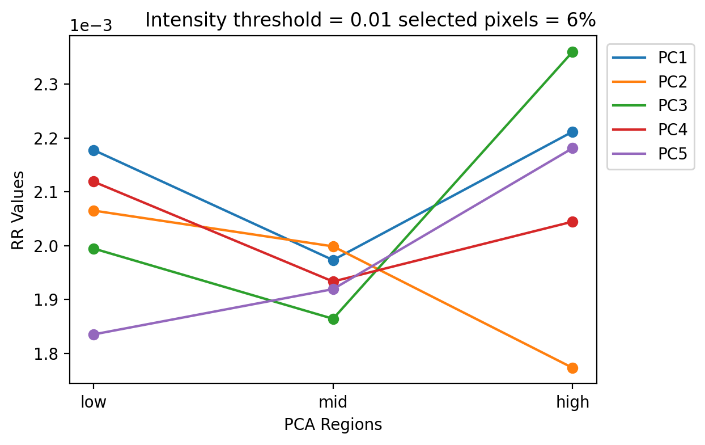

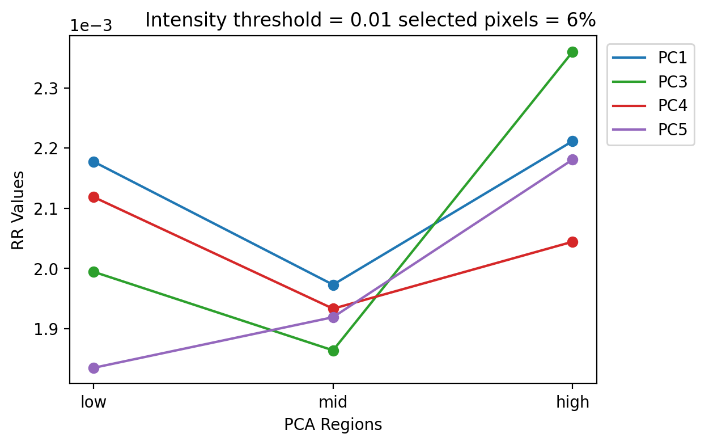

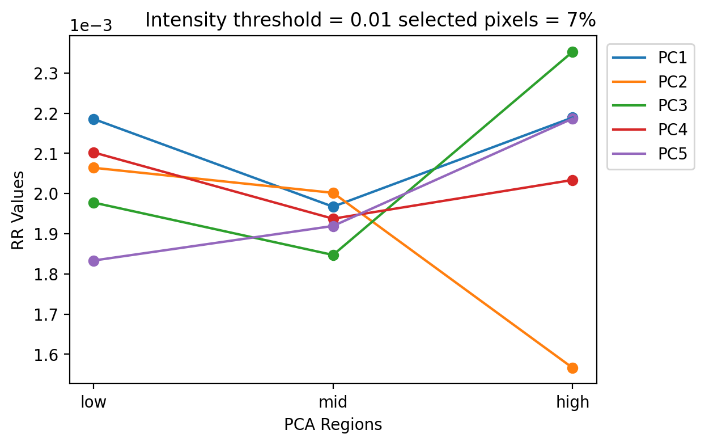

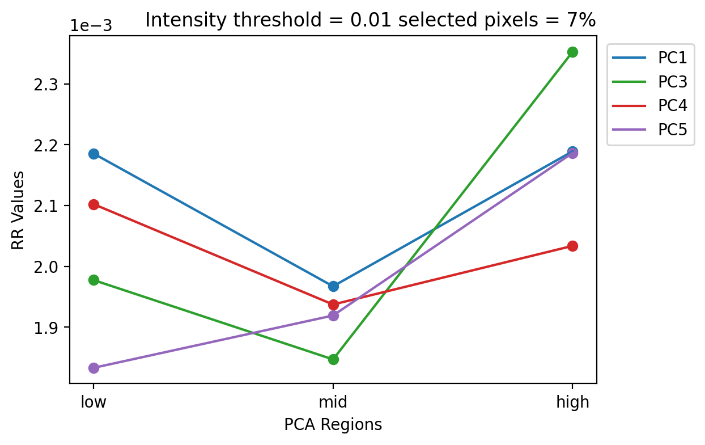

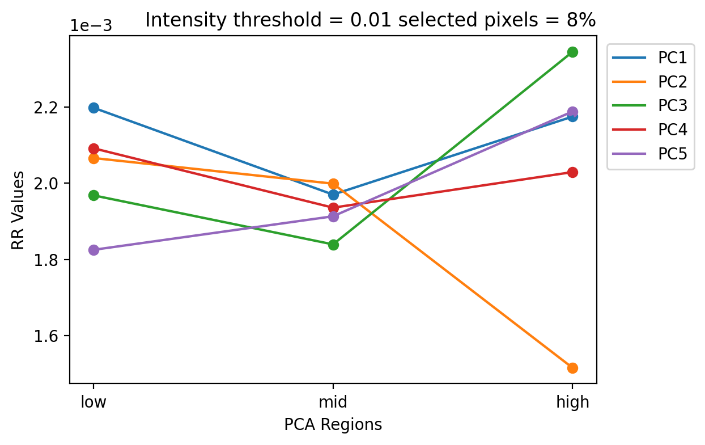

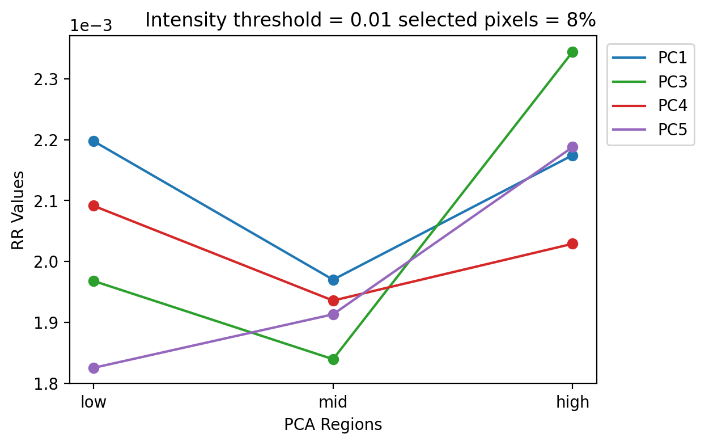

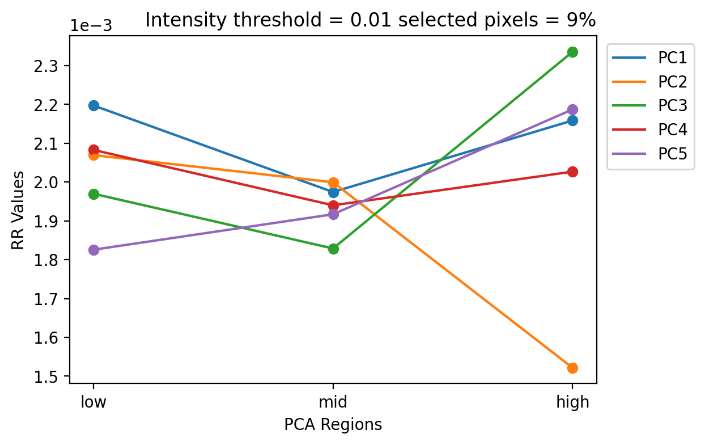

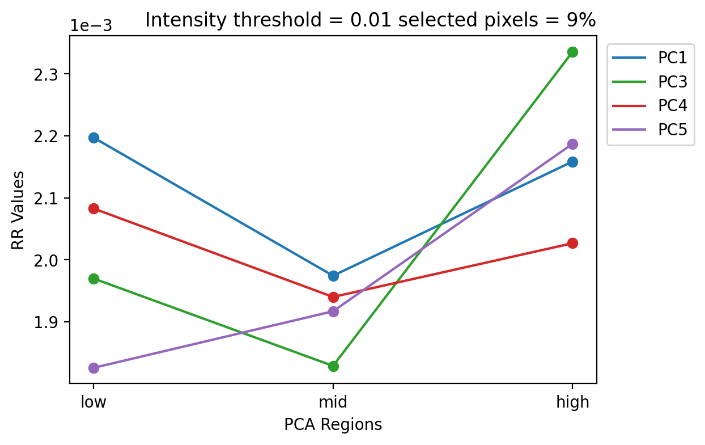

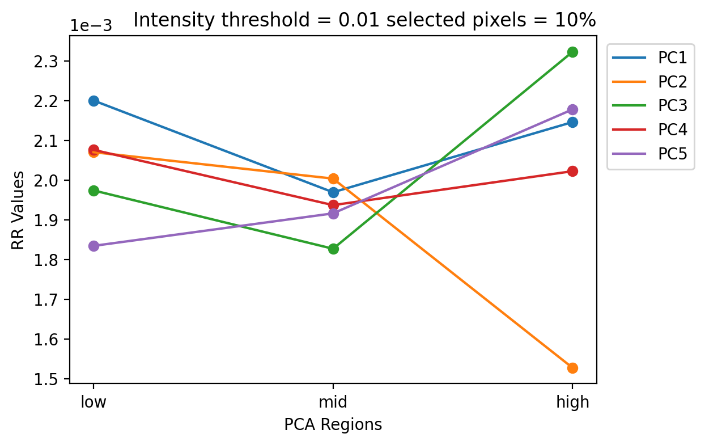

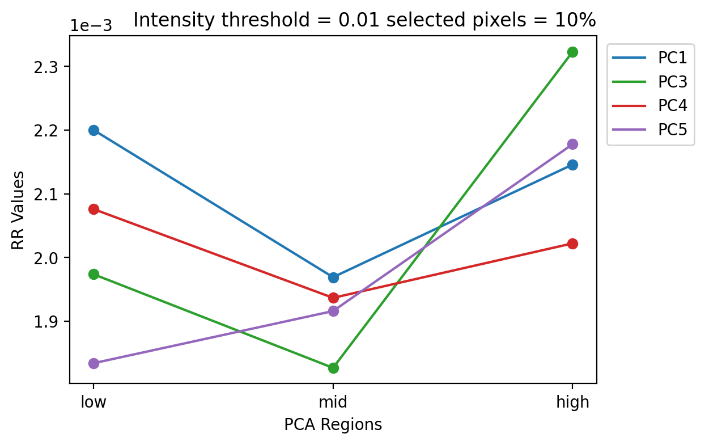

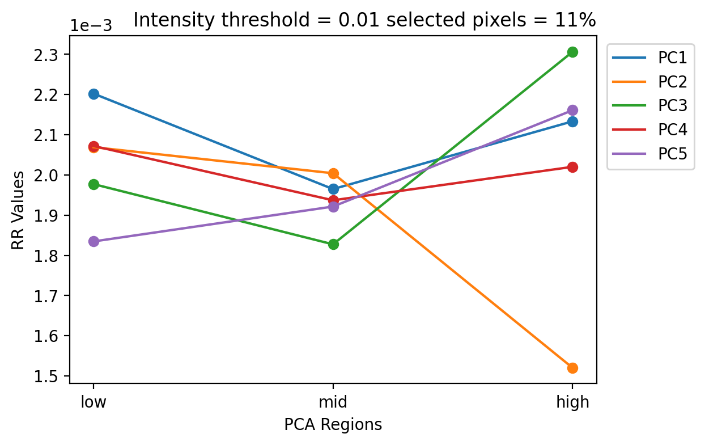

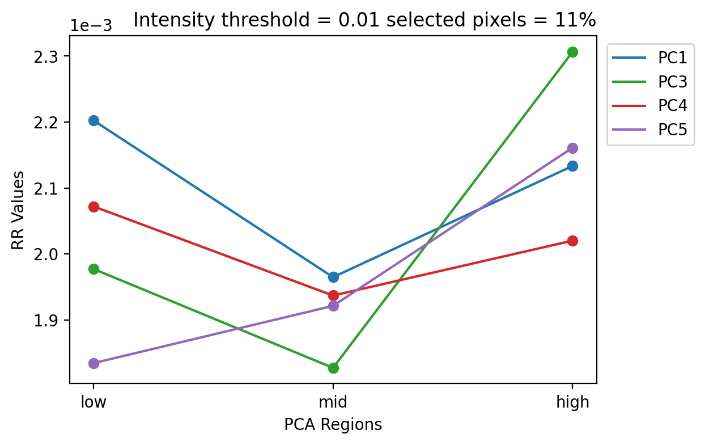

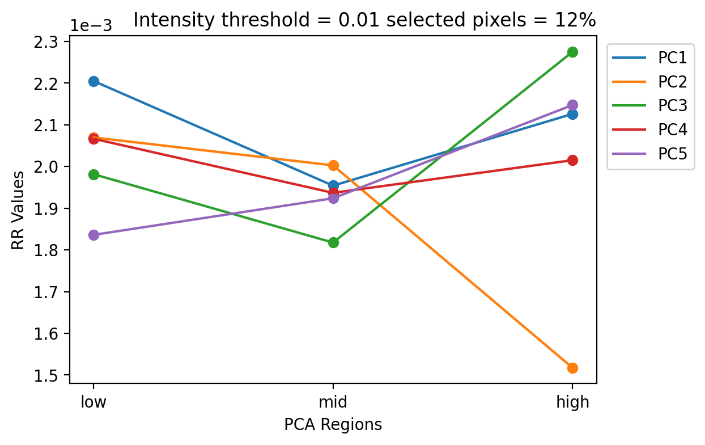

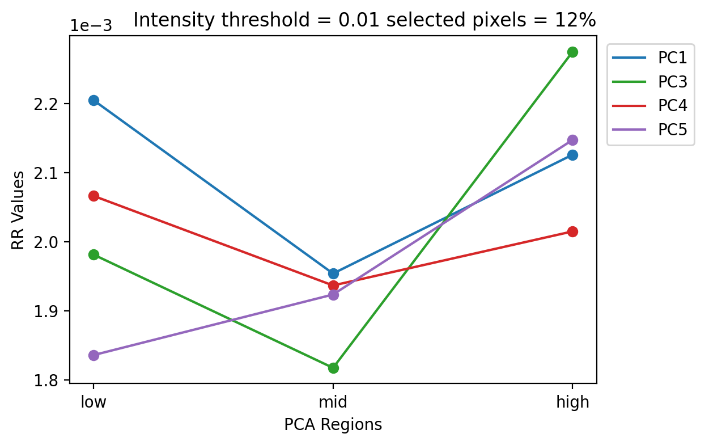

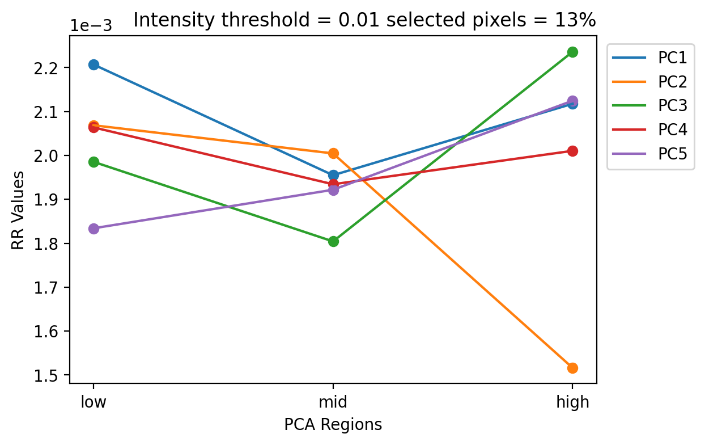

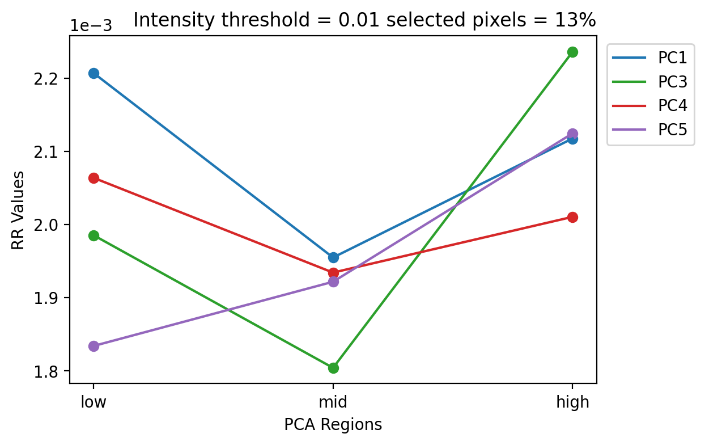

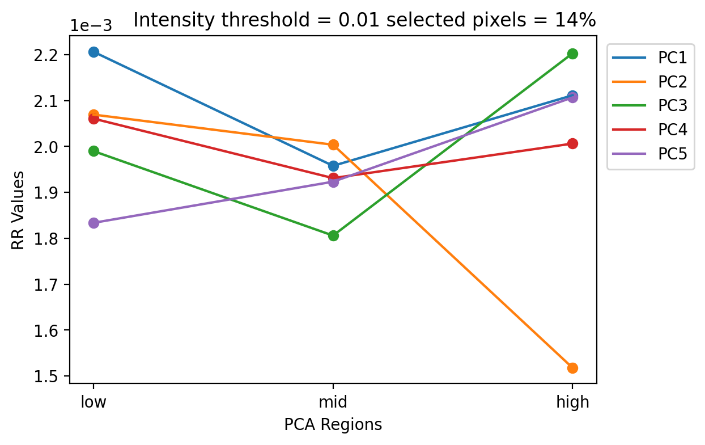

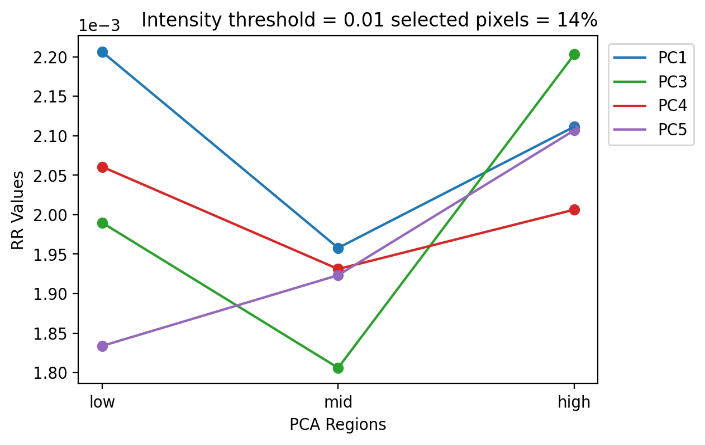

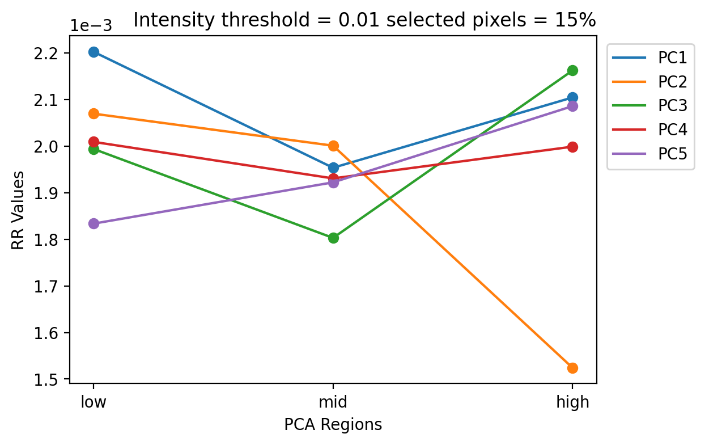

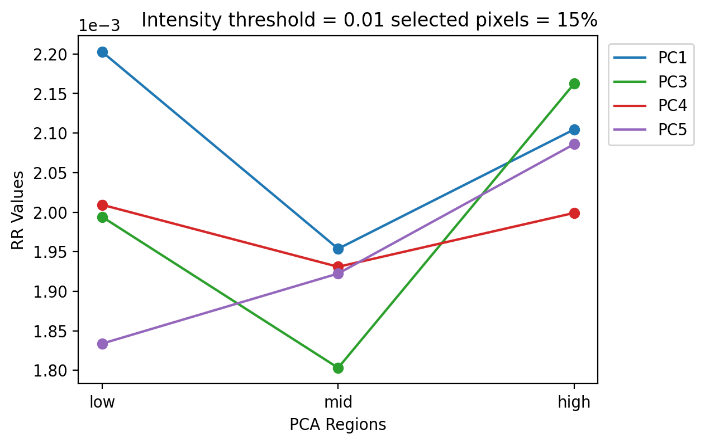

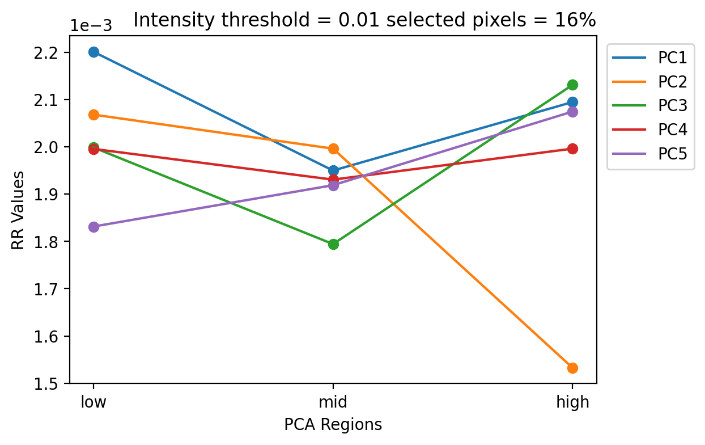

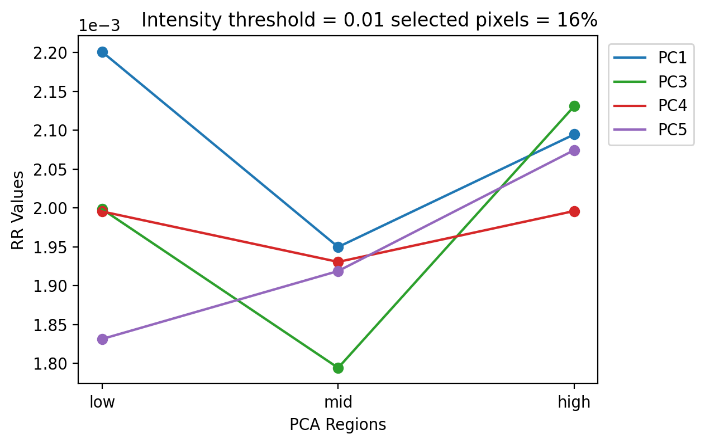

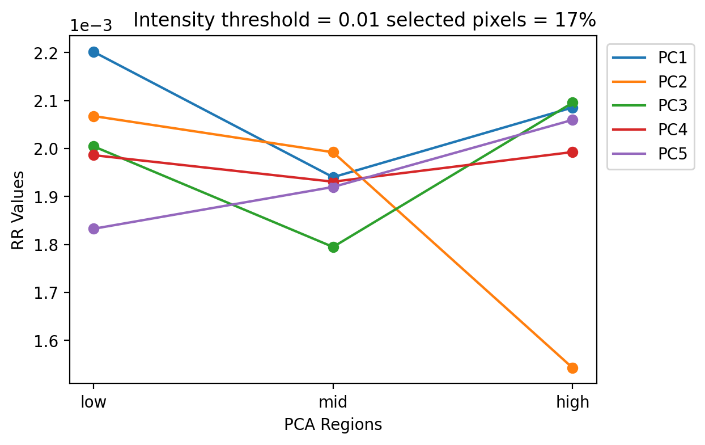

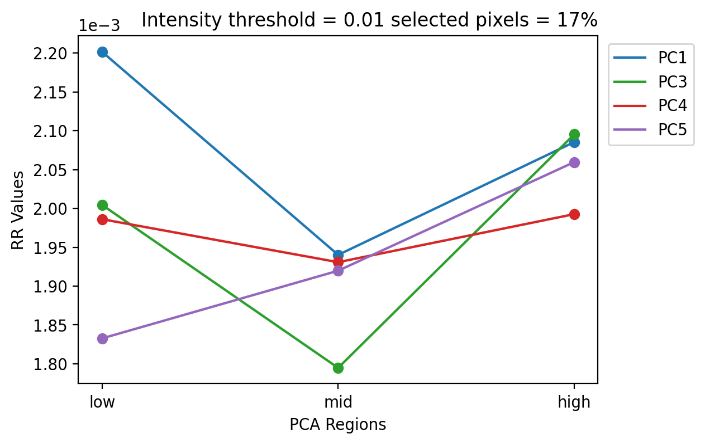

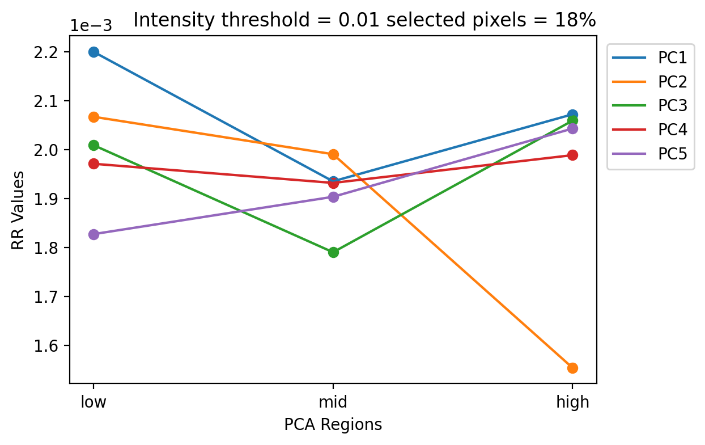

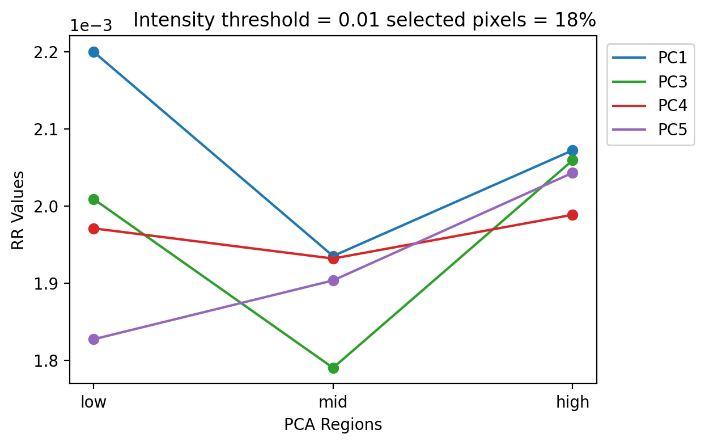

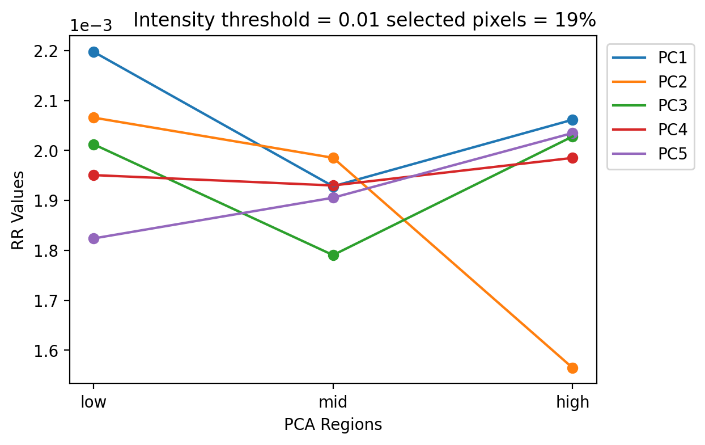

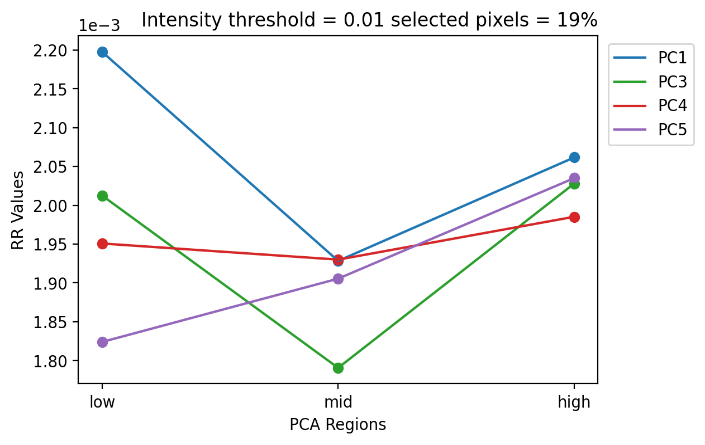

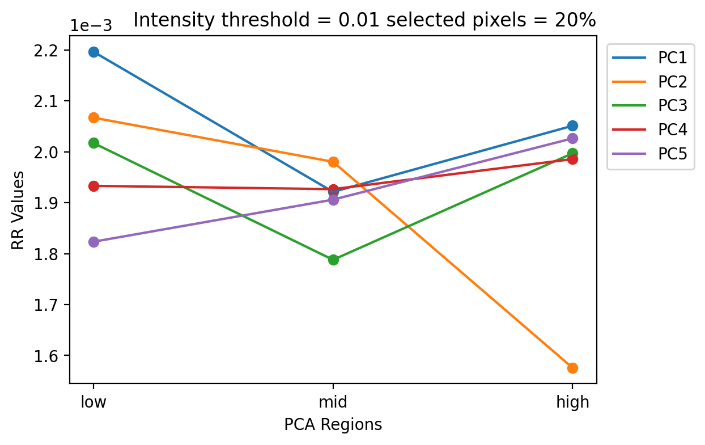

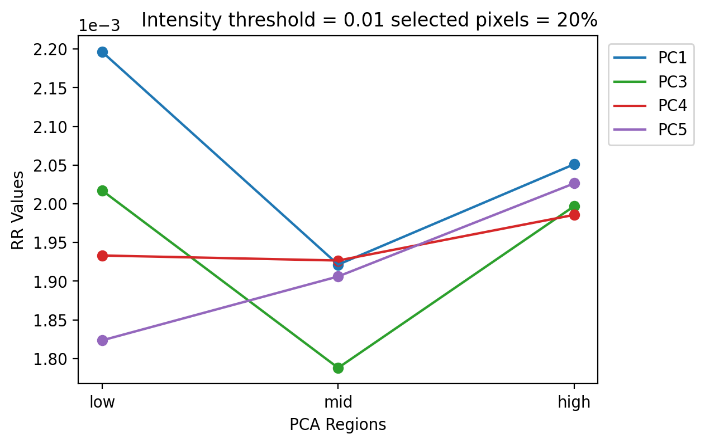

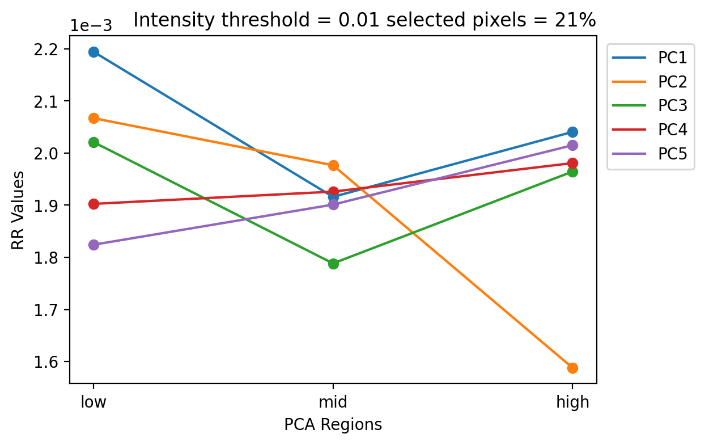

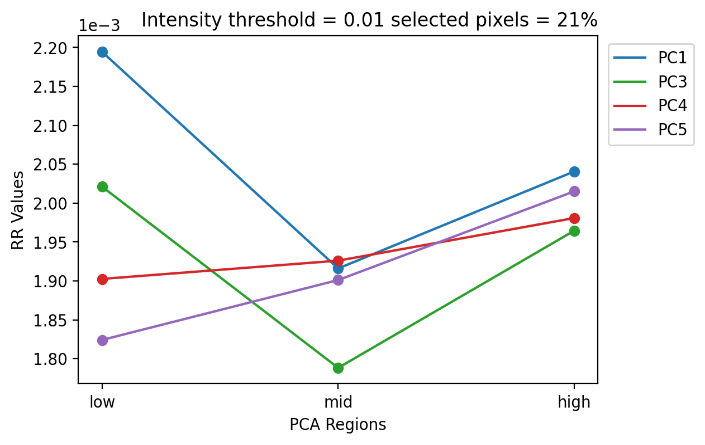

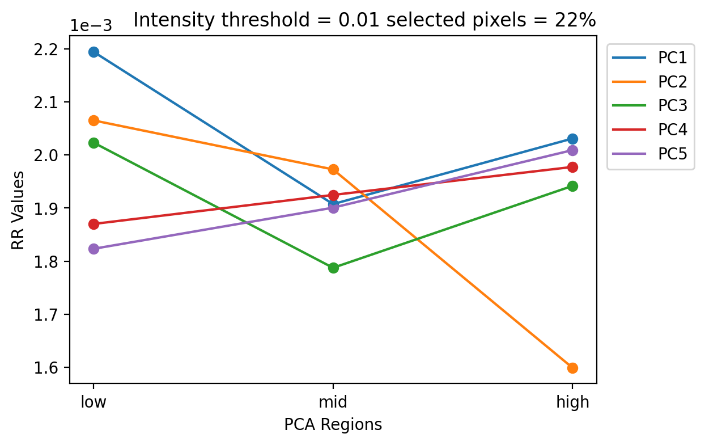

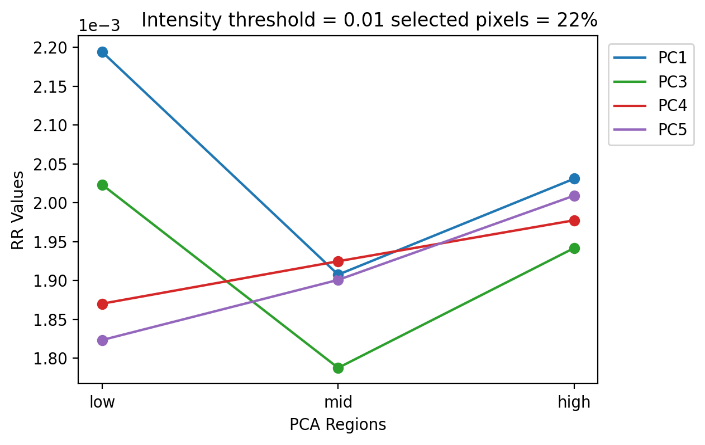

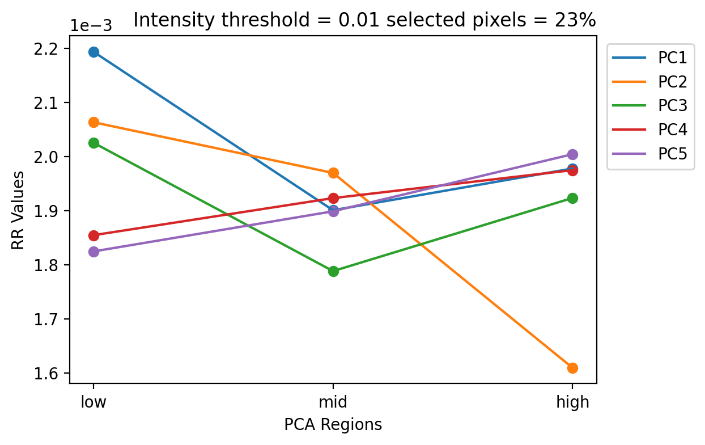

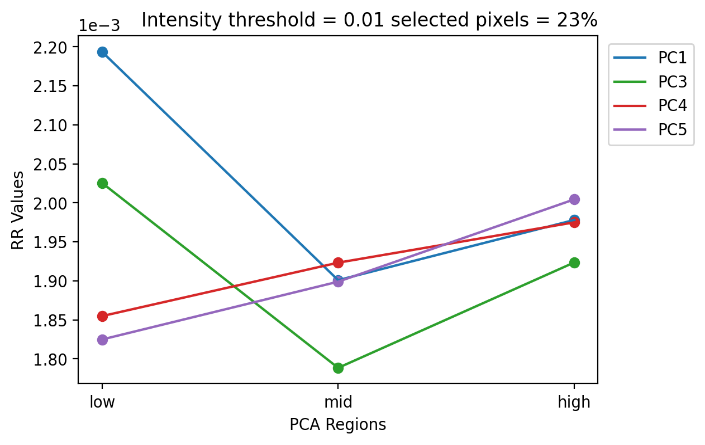

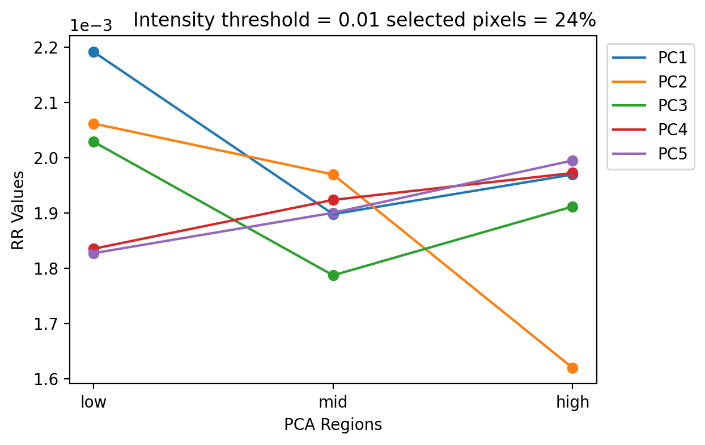

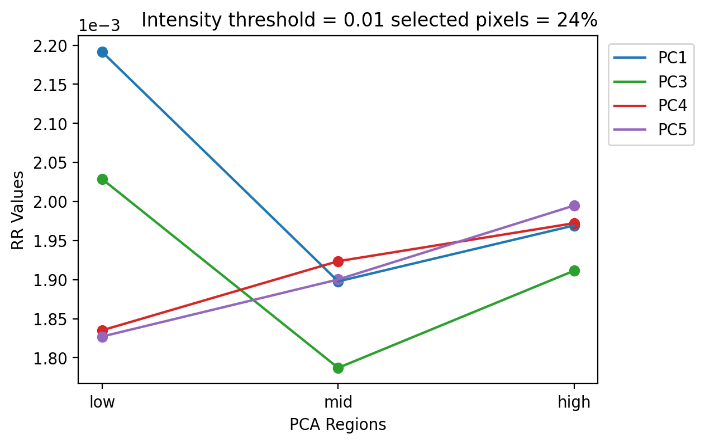

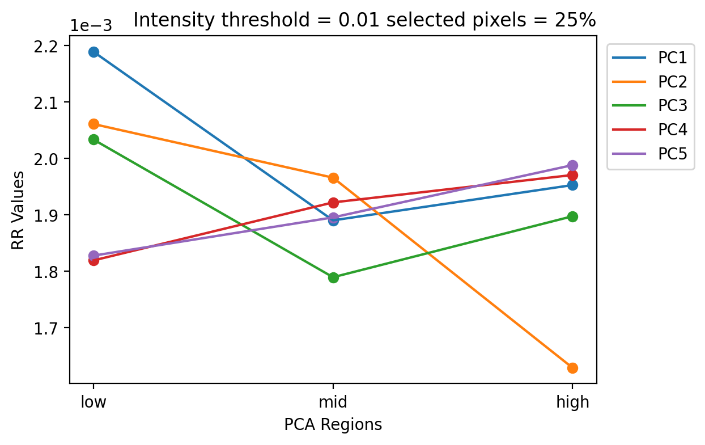

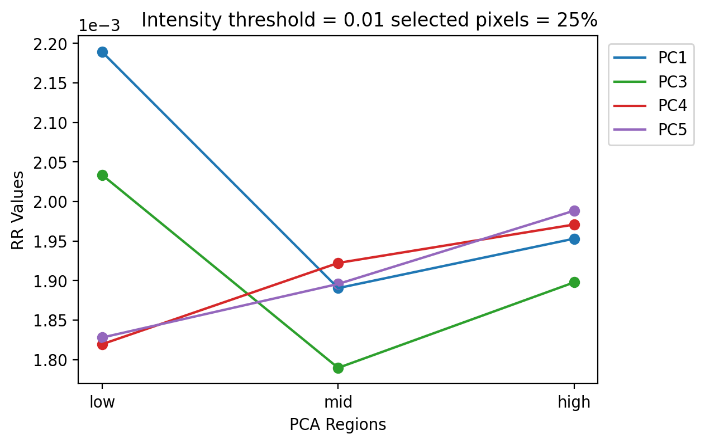

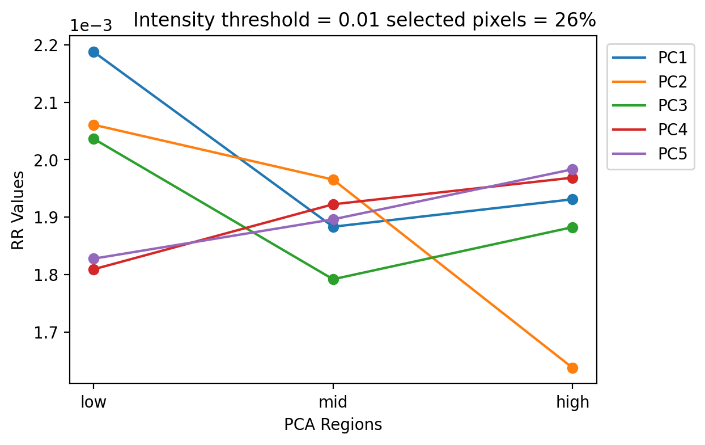

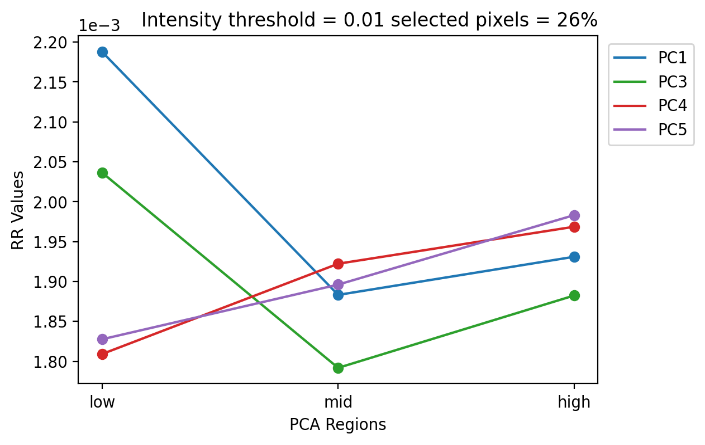

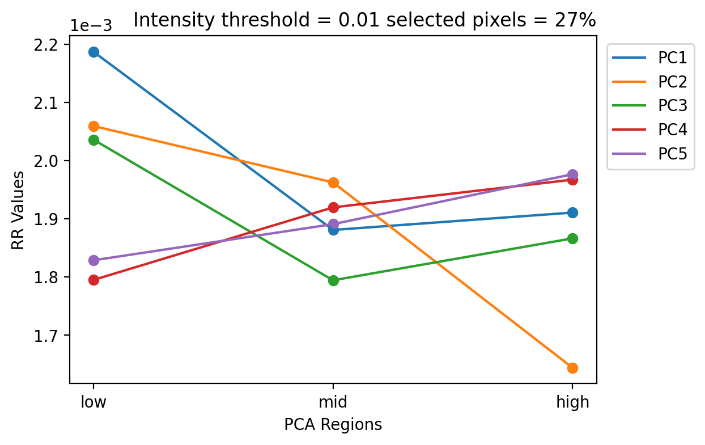

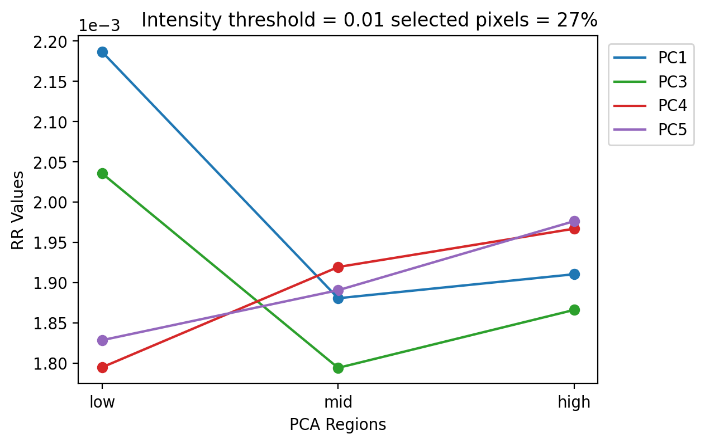

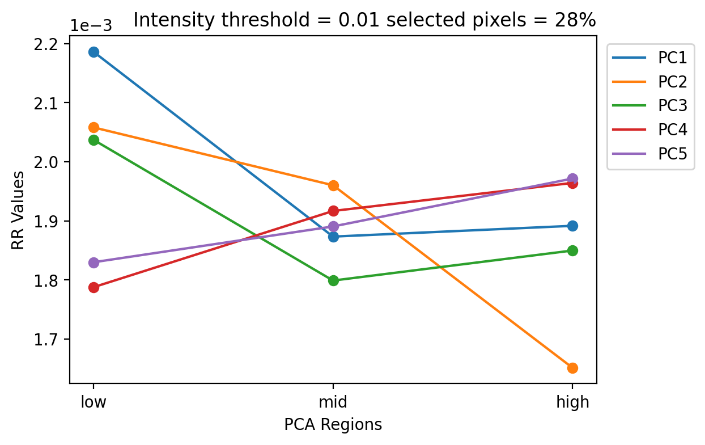

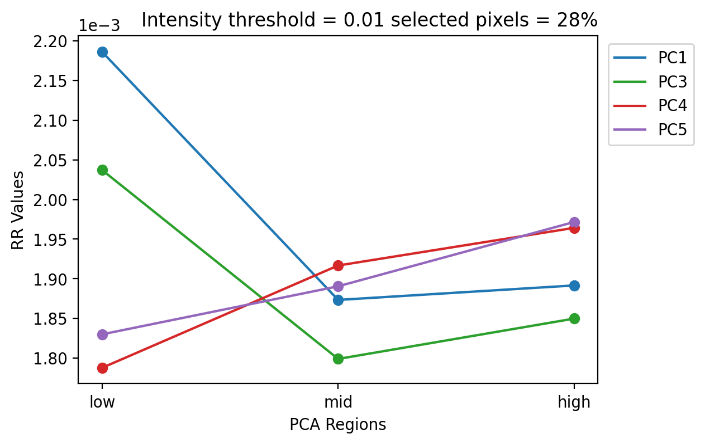

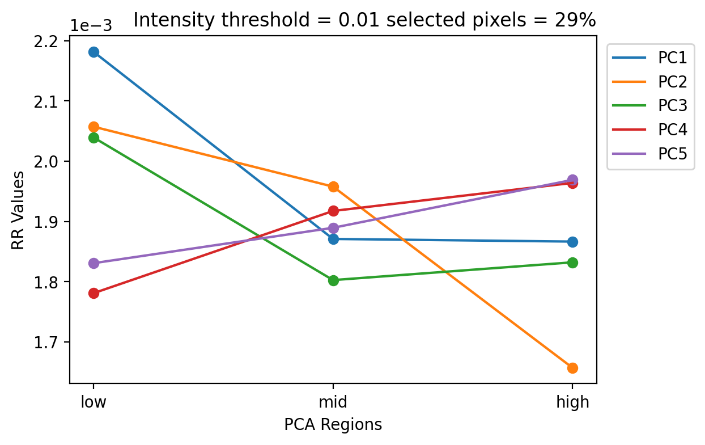

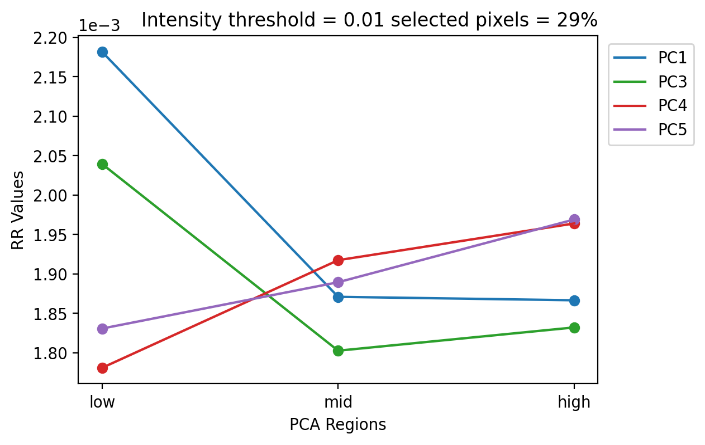

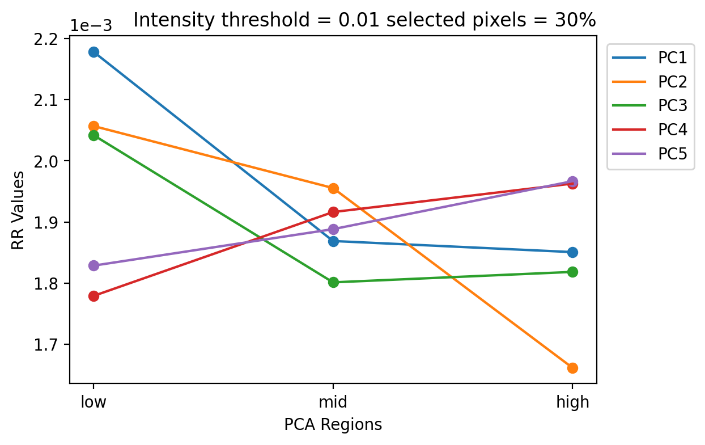

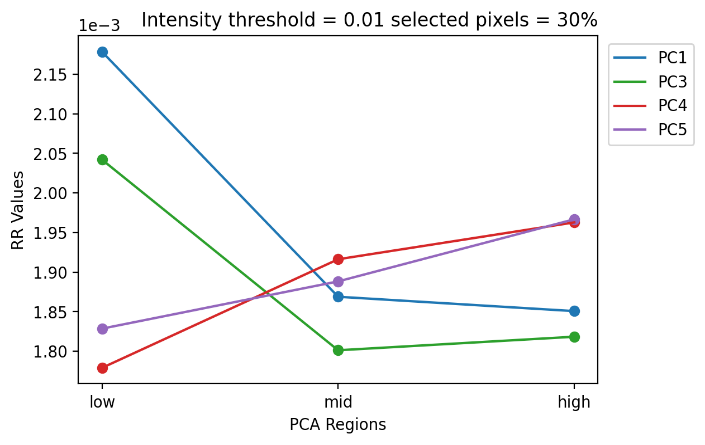

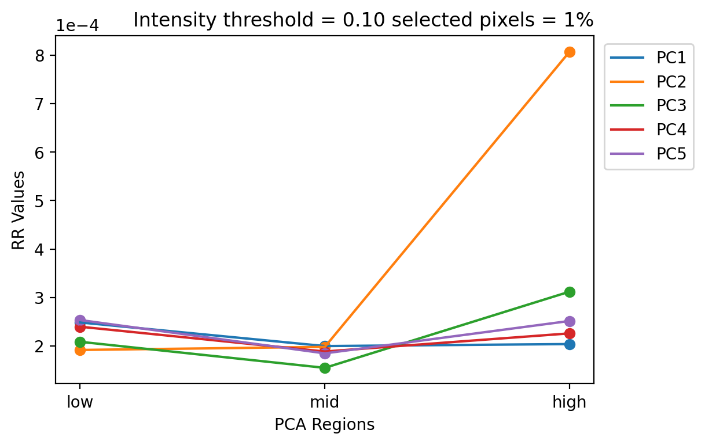

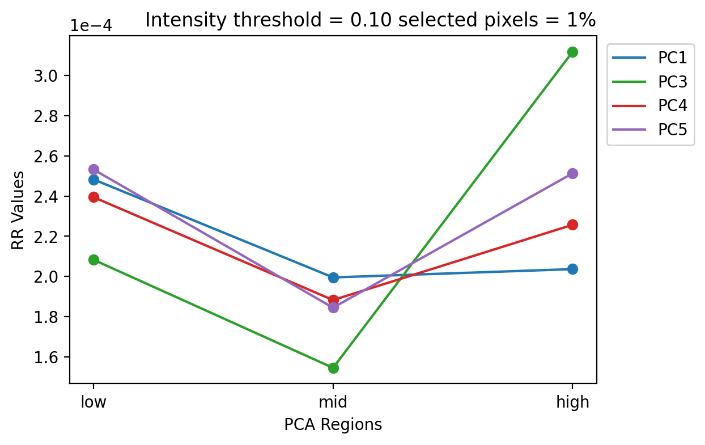

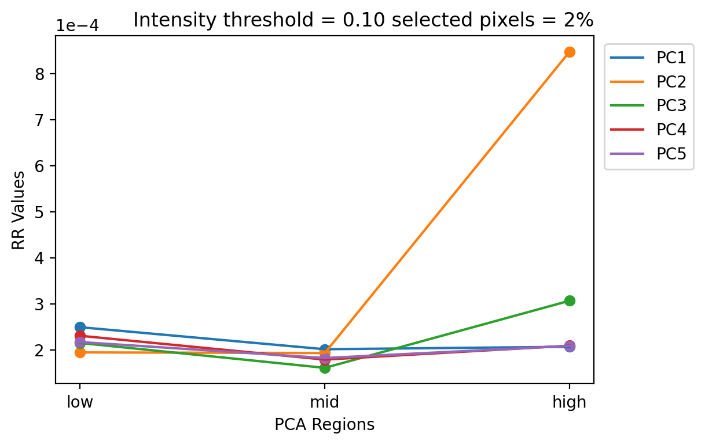

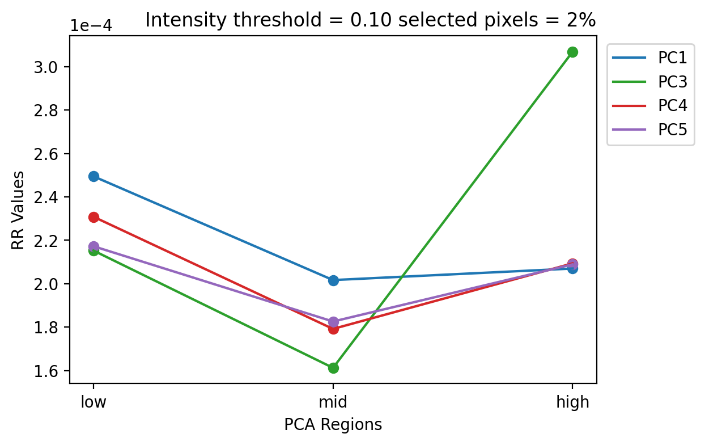

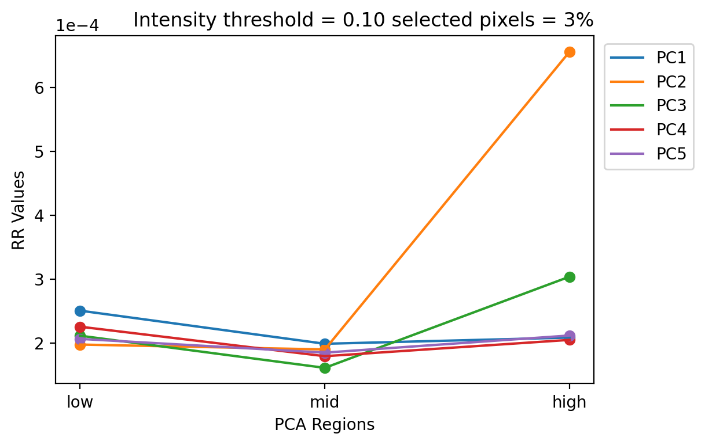

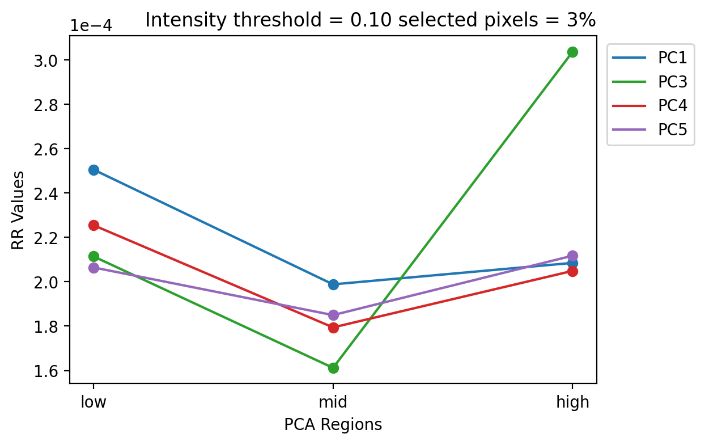

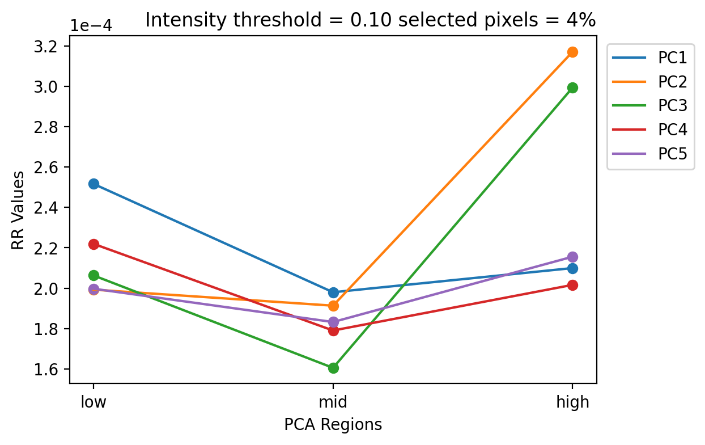

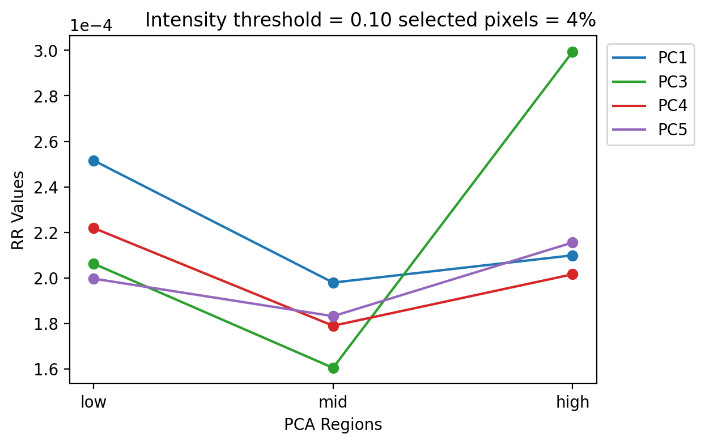

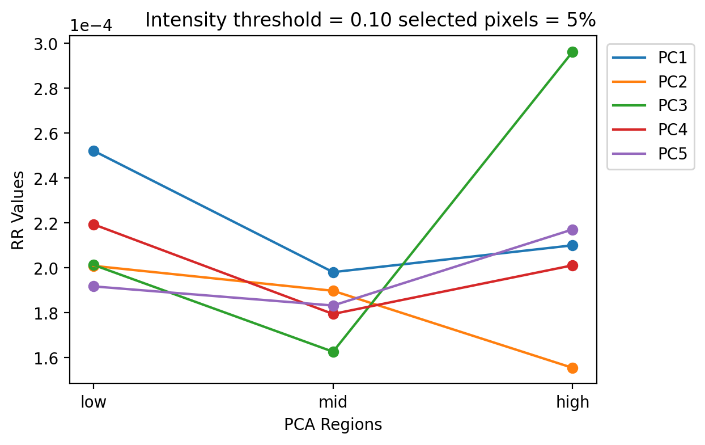

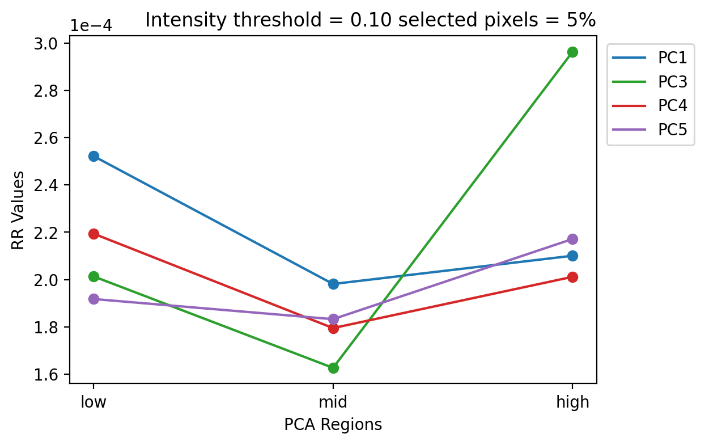

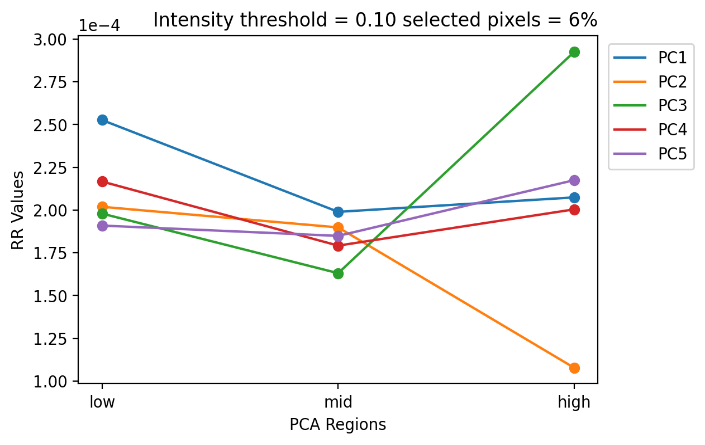

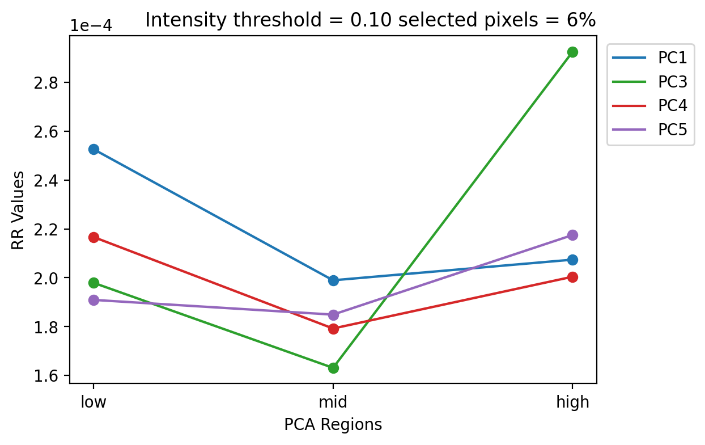

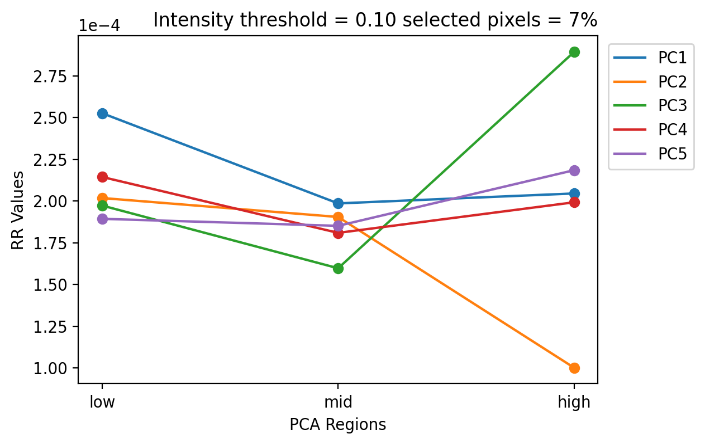

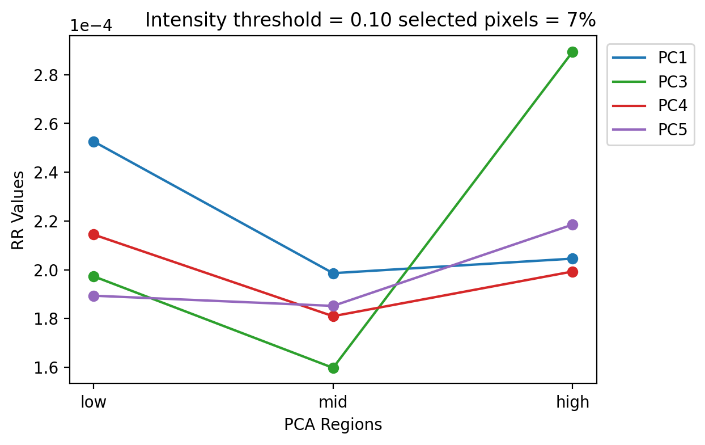

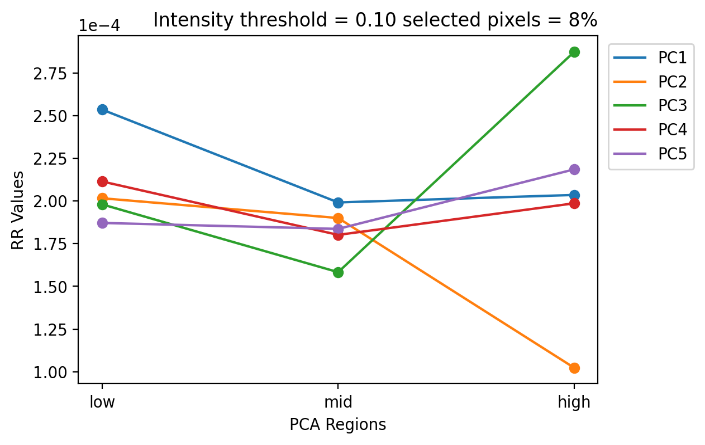

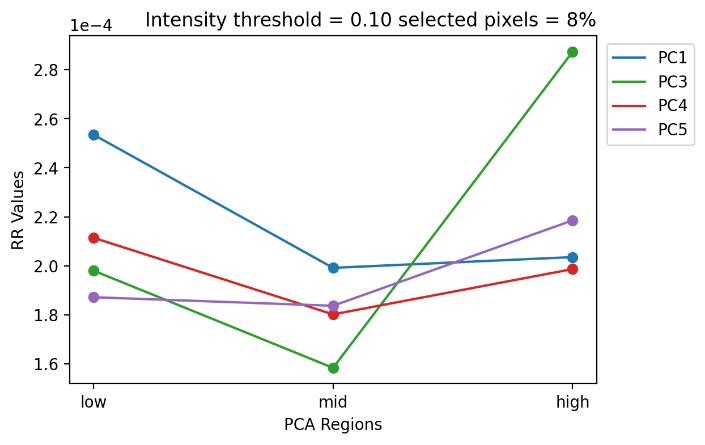

2-D V plots for global normalization with root mean squared normalized results

RMS pre-processing for PCA was used for pixel selection. Global normalization with RMS pre-processing gives comparable distinctions of similarity to the RMS normalized intensities with local normalization. The local method was still determined to be better since the V shape of PC 1 is present throughout the first 10% of selected pixels which makes up the strongest correlated pixels.

2-D V plots for local normalization with non-normalized PCA results

Non-normalized PCA results were used for pixel selection. Local normalization without RMS pre-processing at first showed promising results with good V shapes in the first few percentages of pixels selected. However for intensity threshold 0.01 and 0.10, after 5% the V shape completely disappears for all PCs.

2-D V plots for global normalization with non-normalized PCA results

Non-normalized PCA results were used for pixel selection and no RMS pre-processing for global normalization. The method only showed one V plot for PC 4 that disappears after 3% selected pixels.

2-D V plots for localTIC normalization with non-normalized PCA results

Non-normalized PCA results were used for pixel selection and no RMS pre-processing for global normalization. The method only showed one V plot for PC 4 that disappears after 3% selected pixels. This shows how even though PCA could find biological regions, the workflow can appear to fail simply due to a bad normalization method.

Russel-Rao 3-D V Plots

Optimal set of parameters based on the E-index calculations. A-E correspond to PC 1-5, respectively. All PCs follow the expected trend of decreasing similarity as the coincidence threshold increases. Intensity threshold was set to 0.1 and the percent selected pixels was 1%.

Medoid Spectra

Local normalization with 1% selected pixels, Intensity threshold 0.10, and RMS normalizes PCA results

PC 2

PC 3

PC 4

PC 5

Local normalization with 1% selected pixels, Intensity threshold 0.01, and RMS normalizes PCA results

Significantly fewer peaks are correlated to the PCA loadings due to the intensity threshold being lower than the range of variations for most of the selected PCA peaks.

PC 1

PC 2

PC 3

PC 4

PC 5

Mouse brain image A

Imaging conditions

Ten micrometer thick transverse mouse brain sections were prepared using CM 3050S cryostat (Leica Biosystems, Vista, CA) and stored in a -80°C freezer. The tissue sections were placed in the desiccator for 30 min prior to MALDI matrix application. A 2,5-Dihydroxybenzoic acid (DHB) MALDI matrix layer was applied using a home-built sublimation apparatus. The mouse brain sections were then stored in the desiccator for another 30 min before matrix-assisted laser desorption/ionization (MALDI) imaging with a 7T solariX Fourier transform ion cyclotron resonance mass spectrometer (FTICR) (Bruker Daltonics, Billerica, MA). Imaging parameters were optimized for known lipid profiles^24^ in positive ion mode between *m/z* 400-1000. A laser power of 37% with 750 shots per pixel and a free induced decay (FID) of 0.4893 s were chosen. The resolving power for *m/z* 772.5255 was 34090 based on the full width half maximum (FWHM) mass resolution. Spectra file sizes were set to 256 kB and 98% data reduction was done during acquisition. Spatial resolution and SmartWalk sampling pattern were set to 150 µm and the final image contained 4,211 pixels with a file size of 4.77 GB.

Principal component analysis

PCA was calculated using SCiLS Lab pro by Bruker Daltonics with no normalization, scaling, nor denoising.

Ion images for PCA. From top left to bottom right, *m/z* are as follows: 459.890, 496.351, 554.295, 561.534, 732.560, 732.600, 734.560, 753.594, 756.557, 758.579, 760.570, 769.567, 772.531, 782.557, 798.510, 806.501, 810.578, 834.583, 844.507, 848.548, and 872.542.

Spatial-expression images for PC 1-5.

Pseudo-spectra for PC 1-5 from top to bottom.

Pareto plot for PC 1-5. Explained variance for each PC is as follows: 93.55%, 2.74%, 2.49%, 0.50%, and 0.24%, respectively. The cumulative sum is 93.55%, 96.29%, 98.77%, 99.27%, and 99.51%

E-index Plots

All E-index calculations failed to find optimal set of parameters and did not exhibit any unique trends. The intensity threshold that resulted in the largest E value from local normalizationwas selected as the optimal intensity threshold and the selected pixel percentage was hand-picked.

local normalization from non-normalized PCA results

global normalization from non-normalized PCA results

Russel-Rao 2-D V Plots

Optimal parameters were found to be intensity threshold of 0.11 with 8% selected pixels and local normalization.

No RMS pre-processing was done.

2-D V plots for local normalization with non-normalized PCA results

2-D V plots for global normalization with non-normalized PCA results

Global normalization, in general, fails to distinguish region similarity with the exception of PC 3.

2-D V plots for globalTIC normalization with non-normalized PCA results

GlobalTIC completely fails to show a distinction of region similarity. This is due to a majority of the data not being above the intensity threshold when normalized to the largest total ion count.

Russel-Rao 3-D V plots for optimal parameters

Optimal parameters: intensity threshold = 0.11, selected pixels percent = 8%, normalization = local.

Mouse brain image B

Imaging conditions

Ten micrometer thick transverse mouse brain sections were prepared using CM 3050S cryostat (Leica Biosystems, Vista, CA) and stored in a -80°C freezer. The tissue sections were placed in the desiccator for 30 min prior to MALDI matrix application. A 2,5-Dihydroxyacetophenone (DHA) MALDI matrix layer was applied using a home-built sublimation apparatus. The mouse brain sections were then stored in the desiccator for another 30 min before matrix-assisted laser desorption/ionization (MALDI) imaging with a 7T solariX Fourier transform ion cyclotron resonance mass spectrometer (FTICR) (Bruker Daltonics, Billerica, MA). Imaging parameters were optimized for known lipid profiles^24^ in positive ion mode between *m/z* 400-2000. A laser power of 26% with 500 shots per pixel was chosen. The resolving power for *m/z* 798.541 was 33000 based on the full width half maximum (FWHM) mass resolution. Spectra file sizes were set to 256 kB and 98% data reduction was done during acquisition. Spatial resolution and SmartWalk sampling pattern were set to 100 µm and the final image contained 10,318 pixels with a file size of 12.0 GB.

Principal component analysis

PCA was calculated using SCiLS Lab pro by Bruker Daltonics with no normalization, scaling, nor denoising.

Ion images of *m/z* values used for PCA. From top left to bottom right *m/z* are as follows: 554.290, 555.292, 558.320, 731.278, 732.608, 734.199, 758.568, 760.584, 772.460, 782.564, 798.542, 806.467, 810.454, 834.455, 844.468, 848.515, and 872.556.

Spatial-expression images for PC 1-5. Although some parts of brain structures can be discerned, they are not strongly expressed by the score values. This shows PCA failed to identify biological regions with strong correlations, largely due to the poor ion images.

Pseudo-spectra for PC 1-5 from top to bottom, respectively.

Pareto plot of explained variance for PC 1-5. Explained variance for PC 1-5 is as follows: 75.79%, 14.23%, 8.322%, 0.5471%, and 0.3682%. Cumulative sum of explained variance for PC 1-5 is as follows: 75.79%, 90.02%, 98.34%, 98.89%, and 99.25%.

E-index Plots

local normalization from non-normalized PCA results

All the E-values resulted in negative values except for when the squared weight function was used (a few points of the *E_m­_wsq_* resulted in negative values but mostly positive). These values were the result of a “Ʌ” shape meaning the mid region was found to have higher similarity than the low and high regions. This is likely due to the PCA failing to strongly correlate biological regions in the mouse brain image. The positive values seen with the weighted squared functions are a result of the inverse sum coefficient and the absolute value function carrying the negative signs. The absolute value function was used to help weed out the negative differences between the low/high regions and the mid, assuming most of the results were positive differences.

Russel-Rao 2-D V Plots

2-D V plots for local normalization with non-normalized PCA results

All tested combinations of normalization methods, intensity threshold, and selected pixel percentages results in the “Ʌ” shape.

Mouse brain image C

Imaging conditions

Ten micrometer thick transverse mouse brain sections were prepared using CM 3050S cryostat (Leica Biosystems, Vista, CA) and stored in a -80°C freezer. The tissue sections were placed in the desiccator for 30 min prior to MALDI matrix application. A 2,5-Dihydroxyacetophenone (DHA) MALDI matrix layer was applied using a home-built sublimation apparatus. The mouse brain sections were then stored in the desiccator for another 30 min before matrix-assisted laser desorption/ionization (MALDI) imaging with a 7T solariX Fourier transform ion cyclotron resonance mass spectrometer (FTICR) (Bruker Daltonics, Billerica, MA). Imaging parameters were optimized for known lipid profiles^24^ in positive ion mode between *m/z* 400-2000. A laser power of 20% with 50 shots per pixel was chosen. The resolving power for *m/z* 760.584 was 39520 based on the full width half maximum (FWHM) mass resolution. Spectra file sizes were set to 256 kB and 98% data reduction was done during acquisition. Spatial resolution and SmartWalk sampling pattern were set to 100 µm and the final image contained 6,763 pixels with a file size of 7.75 GB.

Principal component analysis

PCA was calculated using SCiLS Lab pro by Bruker Daltonics with no normalization, scaling, nor denoising.

Ion images for *m/z* used in PCA. From top left to bottom right the *m/z* are as follows: 456.132, 473.195, 555.922, 731.278, 734.199, 758.568, 760.584, 772.460, 782.564, 798.542, 806.467, 810.454, 834.455, 844.468, 848.515, and 872.556.

Spatial-expression images for PC 1-5.

Pseudo-spectra for PC 1-5 from top to bottom, respectively.

Pareto plot for the explained variance of each PC. Explained variance for PC 1-5 are 87.97%, 4.867%, 2.782%, 1.508%, and 1.286%, respectively. The cumulative sum of the explained variance is as follows: 87.97%, 92.84%, 95.62%, 97.13%, and 98.41%.

E-index Plots

local normalization from non-normalized PCA results

E-index results for local normalization from non-normalized PCA. Both functions of the E-index and all three types of weighted functions result in a local maximum tend. The local maximum is typically with 5-10% selected pixels. The largest local maximum from the *E_m_wsq_* (intensity threshold = 0.17 and selected pixel percent = 7) resulted in the optimal set of parameters for calculation.

The coincidence threshold was tested to see its effect on the E-index. Originally 5% increments from [*n*mod2, *n*-1] was chosen. As the increments are shortened the local maximum becomes more apparent, potentially indicating a better estimate of the optimal parameters when more coincidence thresholds are used. Local maximum for 5% increments was found to be at 6% and the local maximum for 1% increments was found to be 7%.

Russel-Rao 2-D V Plots

2-D V plots for local normalization with non-normalized PCA results

2D V plot of optimal parameters from the E-index calculations.

Russel-Rao 3-D V Plots

3-D V plots for local normalization with non-normalized PCA results

3D V plots for optimal parameters intensity threshold of 0.17 and selected pixels percentage of 7%.
